# The RNA-binding protein RbpB is a central regulator of polysaccharide utilization in gut *Bacteroides*

**DOI:** 10.1101/2023.12.22.572801

**Authors:** Ann-Sophie Rüttiger, Daniel Ryan, Luisella Spiga, Vanessa Lamm-Schmidt, Gianluca Prezza, Sarah Reichardt, Lars Barquist, Franziska Faber, Wenhan Zhu, Alexander J. Westermann

## Abstract

Paramount to human health, symbiotic bacteria in the gastrointestinal tract rely on the breakdown of complex polysaccharides to thrive in this sugar-deprived environment. Gut *Bacteroides* are metabolic generalists and deploy dozens of polysaccharide utilization loci (PULs) to forage diverse dietary and host-derived glycans. The expression of the multi-protein PUL complexes is tightly regulated at the transcriptional level. However, how PULs are orchestrated at translational level in response to the fluctuating levels of their cognate substrates is unknown. Here, we identify the RNA-binding protein RbpB and a family of noncoding RNAs as key players in post-transcriptional PUL regulation. Ablation of RbpB in *Bacteroides thetaiotaomicron* displays compromised colonization in the mouse gut in a host diet-dependent manner. Current dogma holds that individual PULs are regulated by dedicated transcriptional regulators. We demonstrate that RbpB acts as a global RNA binder that directly interacts with several hundred cellular transcripts. This includes a paralogous noncoding RNA family comprised of 14 members, the FopS (family <u>o</u>f <u>p</u>aralogous <u>s</u>RNAs) cluster. Through a series of *in*-*vitro* and *in*-*vivo* assays, we reveal that FopS sRNAs repress the translation of a SusC-like glycan transporter when substrates are limited—an effect antagonized by RbpB. Together, this study implicates RNA-coordinated metabolic control as an important, yet previously overlooked, factor contributing to the *in*-*vivo* fitness of predominant microbiota species in dynamic nutrient landscapes.

## INTRODUCTION

The obligate anaerobic, Gram-negative *Bacteroidota* represents a dominant phylum of the gut microbiota ^1,2^ and influences human health in various ways ^3^. The success of *Bacteroides* spp. in colonizing the mammalian bowel is largely due to their immense metabolic capacities encoded on ∼100 distinct polysaccharide utilization loci (PULs) ^4^. PULs enable these bacteria to catabolize complex dietary polysaccharides and host mucus-derived glycans ^5^. Each PUL encodes sets of membrane-spanning, glycolytic multi-protein complexes, which bind the respective substrate at the cell surface and cleave it into oligosaccharides. These oligosaccharides are imported into the periplasm through TonB-dependent transporters of the SusC family ^6^, and further processed into simple sugars. As individual PULs are substrate-specific, their expression is tightly controlled and responds to the nutritional fluctuations that are commonly associated with the dynamic gut environment. Therefore, PULs include dedicated regulatory factors that spur transcription of their specific PUL operons when the corresponding carbon sources are sensed. This includes SusR-like regulators ^7^ and hybrid two-component systems ^8,9^ that combine both sugar-sensing and gene regulatory functions in a single polypeptide. Alternatively, transcription of PUL systems specific to the processing of host-derived glycans is typically governed by extracytoplasmic function sigma/anti-sigma factor pairs ^10^. However, if and how *Bacteroides* prevent ongoing translation of pre-existing PUL mRNAs once the inducing stimulus fades, is not currently known.

Only recently did RNA-seq studies from us ^11,12^ and others ^13^ map the transcriptome of *Bacteroides* spp. at single-nucleotide resolution. This entailed the identification of hundreds of noncoding RNA candidates, including the small noncoding RNAs (sRNAs) DonS and GibS. DonS is divergently encoded to an *N*-glycan-specific PUL and was shown to repress the expression of this PUL in *Bacteroides fragilis* ^13^, albeit through an unknown mechanism. The conserved sRNA GibS is induced in the presence of *N*-glycans, and binds and represses a glycoside hydrolase-encoding mRNA in *Bacteroides thetaiotaomicron* ^11^. Given these examples and the general importance of RNA-based regulation, it is likely that *Bacteroides* employ much more complex sRNA-based post-transcriptional regulatory networks to optimize fitness in response to dynamic nutrient levels. However, their identification in these bacterial taxa that lack homologs of classical RNA chaperones has proven challenging.

The present study reports the identification of an *in*-*vivo* phenotype, the RNA interactome, a metabolism-associated function, and the underlying molecular mechanism of the first global RNA-binding protein (RBP) in *Bacteroides*. Specifically, we found that a *B*. *thetaiotaomicron* mutant devoid of the RNA recognition motif 1 (RRM-1)-containing protein RbpB ^14,15^ failed to efficiently colonize the mammalian intestine in a diet-dependent manner. Cross-linking immunoprecipitation and sequencing (CLIP-seq) of *B*. *thetaiotaomicron* RbpB demonstrated that this protein is a global RNA binder. Analysis of the RbpB interactome led to the discovery of a complex RNA network, with a highly conserved multicopy sRNA family at its center: the ‘FopS’ cluster (for ‘family <u>o</u>f <u>p</u>aralogous <u>s</u>RNAs’). While the 14 FopS members of *B*. *thetaiotaomicron* exhibit partial sequence similarity to the previously characterized GibS sRNA ^11^, we provide evidence of functional diversification between GibS and FopS. The FopS sRNAs function as post-transcriptional repressors of specific PUL operons by binding to the vicinity of the start codon of the corresponding *susC* homologue, impeding translation of the respective glycan transporter. RbpB counteracts FopS activity, relieving translational repression. Together, this study reports a remarkably complex post-transcriptional control network that allows *Bacteroides* to switch off specific PULs in the absence of the cognate substrate or in the presence of an alternative, preferred carbohydrate source to optimize fitness.

## RESULTS

### RbpB contributes to B. thetaiotaomicron colonization of the mammalian intestine in a diet-dependent manner

*Bacteroides* spp. encode an array of transcription factors that govern carbohydrate utilization in these gut bacteria ^9,10,16–20^. Conversely, there is currently little knowledge as to the extent to which post-transcriptional control impacts on *Bacteroides* metabolic competitiveness. Given that the deletion of the proposed RNA-binder RbpB (BT_1887) in *B*. *thetaiotaomicron* led to the differential expression of several PUL genes ^15^, we assessed the fitness of a *B*. *thetaiotaomicron* Δ*rbpB* mutant in the mammalian gut in response to distinct diets. As conventionally raised mice are resistant to *B*. *thetaiotaomicron* colonization ^21^, we treated C57BL/6 mice with an antibiotic cocktail to promote *B*. *thetaiotaomicron* engraftment. We then inoculated mice fed a conventional diet with an equal ratio of the *B*. *thetaiotaomicron* wild-type strain and an isogenic Δ*rbpB* mutant, and determined the abundance of each strain in cecal and colonic contents six days post-inoculation by plating on selective media (Fig. 1a). Remarkably, the Δ*rbpB* mutant displayed a significant fitness disadvantage compared to the wild-type counterpart in both intestinal sites (Fig. 1b). The magnitude of this attenuation ranged in between the large impact of deleting the transcriptional master regulator of polysaccharide utilization, Cur ^22^, and the relatively mild *in*-*vivo* phenotypes of individual PUL-encoded transcriptional regulators (e.g. ^23,24^). Importantly, the RbpB-associated fitness defects were largely abolished when mice were fed with a fiber-free diet (Fig. 1b). Together, these data underscore the importance of the *Bacteroides* RNA-binding candidate RbpB for carbon source adaptation *in vivo* and prompted us to investigate the cellular function of this protein in more detail.

**Figure 1:**
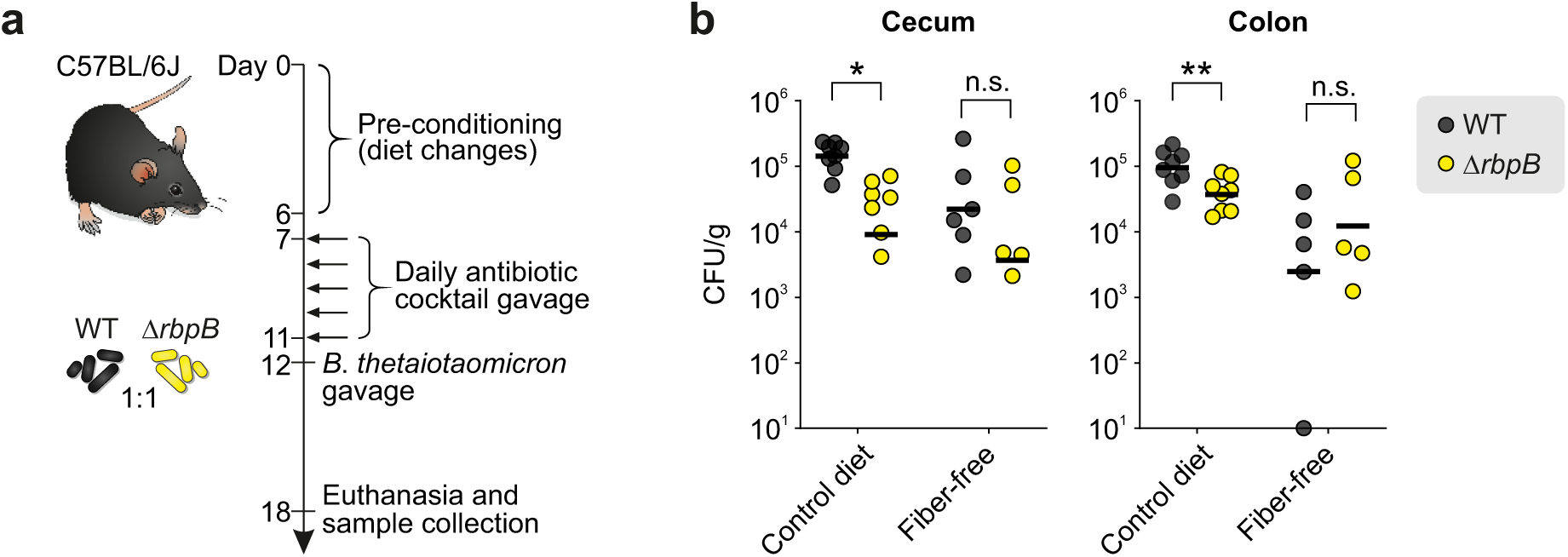
Role of *B*. *thetaiotaomicron* RbpB in the colonization of the mouse gut. **a,** Schematic representation of the experimental outline. One week before antibiotic treatment, C57BL/6J mice were switched to a fiber-free diet or remained on a control diet until the end of the experiment. Antibiotic cocktails were administered by oral gavage daily for 5 days. After antibiotic treatment, mice were inoculated with an equal mixture of each 0.5 × 10^9^ CFU of the *B*. *thetaiotaomicron* wild-type and Δ*rbpB* strain. After 6 days, cecal and colonic tissue was collected and the bacterial numbers determined by selective plating. **b,** The abundance of *B*. *thetaiotaomicron* wild-type (black) or Δ*rbpB* (yellow) in the cecal (left) or colonic (right) contents were determined by selective plating. The data are combined over two independent experiments, each comprised of three biological replicates. Black horizontal lines represent the geometric means. \**p* < 0.05; \*\**p* < 0.01; n.s., not statically significant based on two-tailed Student’s *t*-tests.

### RbpB acts as a global RNA-binding protein in B. thetaiotaomicron

RbpB was reported to possess the ability to bind synthetic single-stranded RNA ‘pentaprobes’ *in vitro* ^15^. However, whether *Bacteroides* RbpB acts as a global RNA binder *in vivo* —as opposed to a test tube—was previously not assessed. To explore the *in*-*vivo* functions of RbpB, we constructed an epitope-tagged version of RbpB and stably integrated it into the chromosome of *B*. *thetaiotaomicron*. *In* -*vivo* crosslinking and immunoprecipitation (CLIP) experiments (Suppl. Fig. S1a) revealed the characteristic crosslink-induced signal on an autoradiograph (Fig. 2a), which was sensitive to RNase I—but not to DNase I—treatment (Suppl. Fig. S1b), suggesting the protein interacts primarily with cellular RNA.

**Figure 2:**
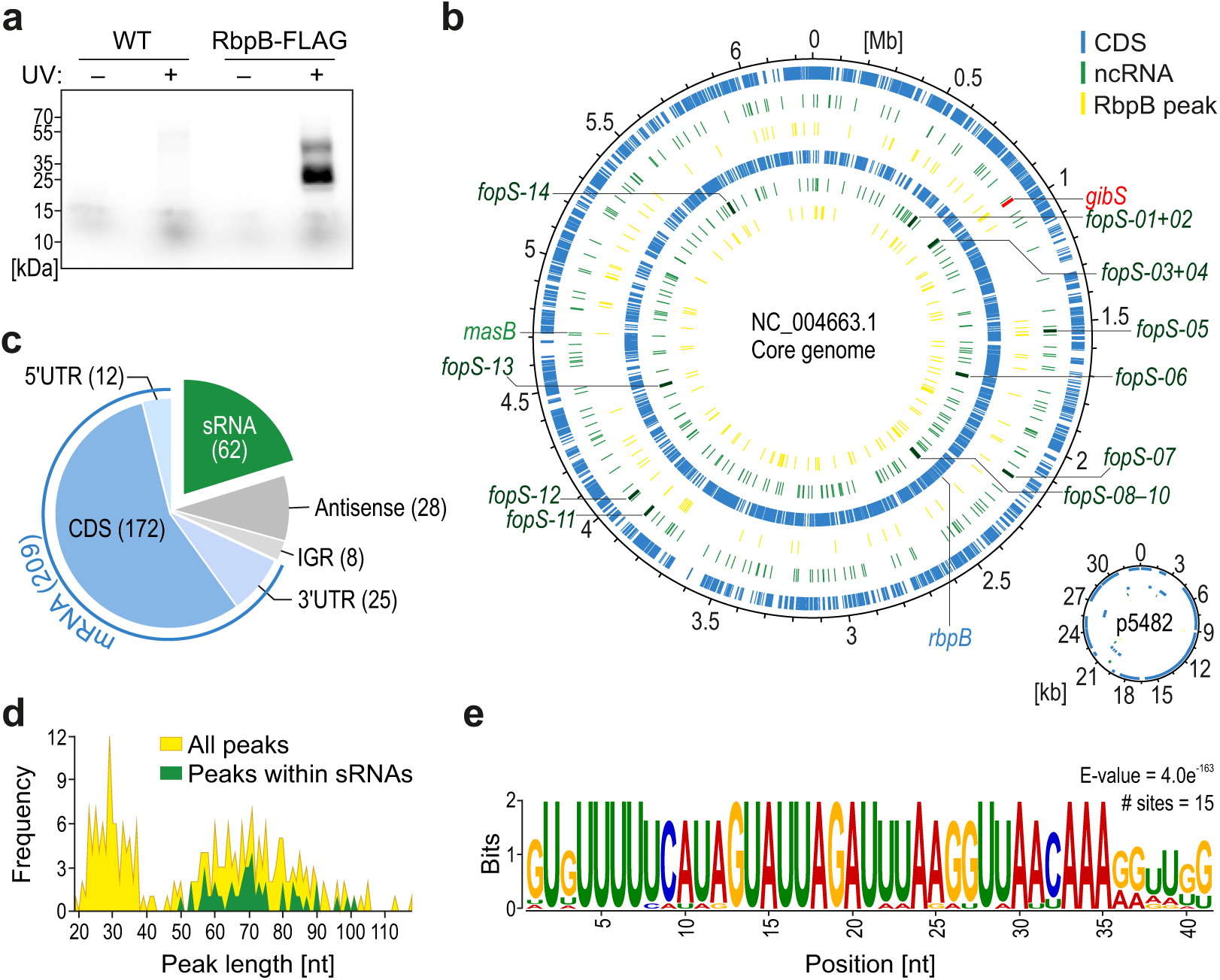
RbpB acts as a global RNA binding protein in *B*. *thetaiotaomicron*. **a,** Autoradiogram of radioactively labeled RNAs covalently bound to RbpB-FLAG after *in*-*vivo* UV crosslinking, immunoprecipitation, and transfer to a nitrocellulose membrane. **b,** Genomic positions of the identified RbpB CLIP peaks in the context of the *B*. *thetaiotaomicron* chromosome and plasmid. The positions of coding sequences (CDS) and noncoding RNA (ncRNA) genes were retrieved from NCBI and published literature ^11^. The *gibS* gene and the 14 FopS-encoding loci (which will become relevant below) are marked. The *rbpB* mRNA itself harbors a RbpB binding site, implying the potential for autoregulation. Inner circles refer to the minus strand and outer circles to the plus strand. **c,** RNA class distribution of RbpB ligands. Numbers in brackets refer to statistically significant and manually confirmed peaks supported by two independent CLIP-seq experiments. Note that the sum of peaks mapping to the different genetic features exceeds the actual total peak number as some peaks mapped to overlapping annotations (such as 3⍰ UTRs and 3’-derived sRNAs). **d,** Peak size distribution. Plotted is the frequencies for individual peak sizes of all unique peaks (yellow) and sRNA peaks (green). **e,** Enriched sequence motif within RbpB ligands.

To map the RbpB interactome at genomic scale, we subjected the co-purified RNA to high-throughput sequencing (CLIP-seq). Two independently performed replicate experiments showed clear read enrichments in crosslinked samples as compared to matched background controls (Suppl. Fig. S1c). We report a total of 285 significantly enriched (*p*_adj_ ≤ 0.05), manually confirmed peaks within 173 different mRNAs (8 peaks in 5’ UTRs, 143 in CDSs, 22 in 3’ UTRs), 62 peaks in 55 different sRNAs (two of which overlap with 3’ UTR peaks and three with 5’ UTR peaks), one or two peaks, respectively, in the 6S and 4.5S housekeeping RNAs, and 14 peaks within intergenic regions (Fig. 2b, c; Suppl. Table 1). Inspecting the peak size distribution revealed two overrepresented footprint lengths (Fig. 2d), reflecting the band pattern on the autoradiogram (Fig. 2a; Suppl. Fig. S1b). While a shorter peak size of ∼30 nt was predominantly associated with mRNAs and intergenic regions, peaks within sRNAs exceeded 50 nt (Fig. 2d). Functional analysis of RbpB-bound mRNAs revealed an enrichment of peaks in transcripts encoding proteins involved in translational processes, such as the biosynthesis of aminoacyl-tRNAs and ribosomal proteins (Suppl. Fig. S2a), implying a translation-related role of this RBP.

In Gram-negative species, global RBPs often stabilize their RNA ligands by shielding them from cellular nucleases ^25–28^. To test for a similar role of *Bacteroides* RbpB, we coupled rifampicin-mediated inhibition of *de*-*novo* transcription to the measurement of RNA decay kinetics ^29^. This analysis suggested that both RbpB depletion and overexpression (Suppl. Fig. S3a), reduce the cellular half-lives of the top-enriched mRNA ligands (Suppl. Fig. S3b), yet hardly of any RbpB-associated sRNAs (Suppl. Fig. S3c). It therefore appears that, rather than merely protecting against degradation, the precise cellular concentration of RbpB defines the stability of its ligands. Closer inspection of CLIP-seq peaks suggested the observed mRNA destabilizations to be independent of the relative position of RbpB binding within those transcripts (Suppl. Fig. S3d). With respect to RbpB-bound noncoding RNAs, drawing from available secondary structure information ^14^, manual inspection revealed the protein to bind preferentially within single-stranded regions (Suppl. Fig. S2b). MEME analysis ^30^ identified several primary sequence motifs enriched in RbpB peaks (Suppl. Fig. S4a), with the most significant comprising a 41 nt-long motif (Fig. 2e). In fact, many of the most strongly enriched sRNA ligands of RbpB contained this sequence around the RBP footprint (Suppl. Fig. S4b). Given this observation, we focused on this sequence motif and the sRNAs containing it.

### A conserved multicopy sRNA family is associated with RbpB

We next set out to determine the prevalence of the above sequence motif. A hidden Markov model-based iterative sequence homology search returned 14 paralogous sequences within the *B*. *thetaiotaomicron* VPI-5482 genome (Fig. 3a). They all fell within annotated sRNAs, which were significantly enriched in the RbpB CLIP-seq dataset. A BLAST search revealed this multicopy sRNA family to be highly conserved within the *Bacteroidota* (Suppl. Fig. S5a), ranging from 12 copies (in *Bacteroides caccae* and *Bacteroides uniformis*) to 15 copies per genome (in *Bacteroides xylanisolvens*) (Suppl. Fig. S5b). Given the presence of the sequence in a Caudovirales phage (Suppl. Fig. S5a)—predicted to target *Bacteroides* based on matching CRISPR spacers ^31^—bacteriophages may have played a role in disseminating the corresponding sRNAs within and across *Bacteroidota* genomes. We term the cluster “family <u>o</u>f <u>p</u>aralogous <u>s</u>RNAs” (FopS).

**Figure 3:**
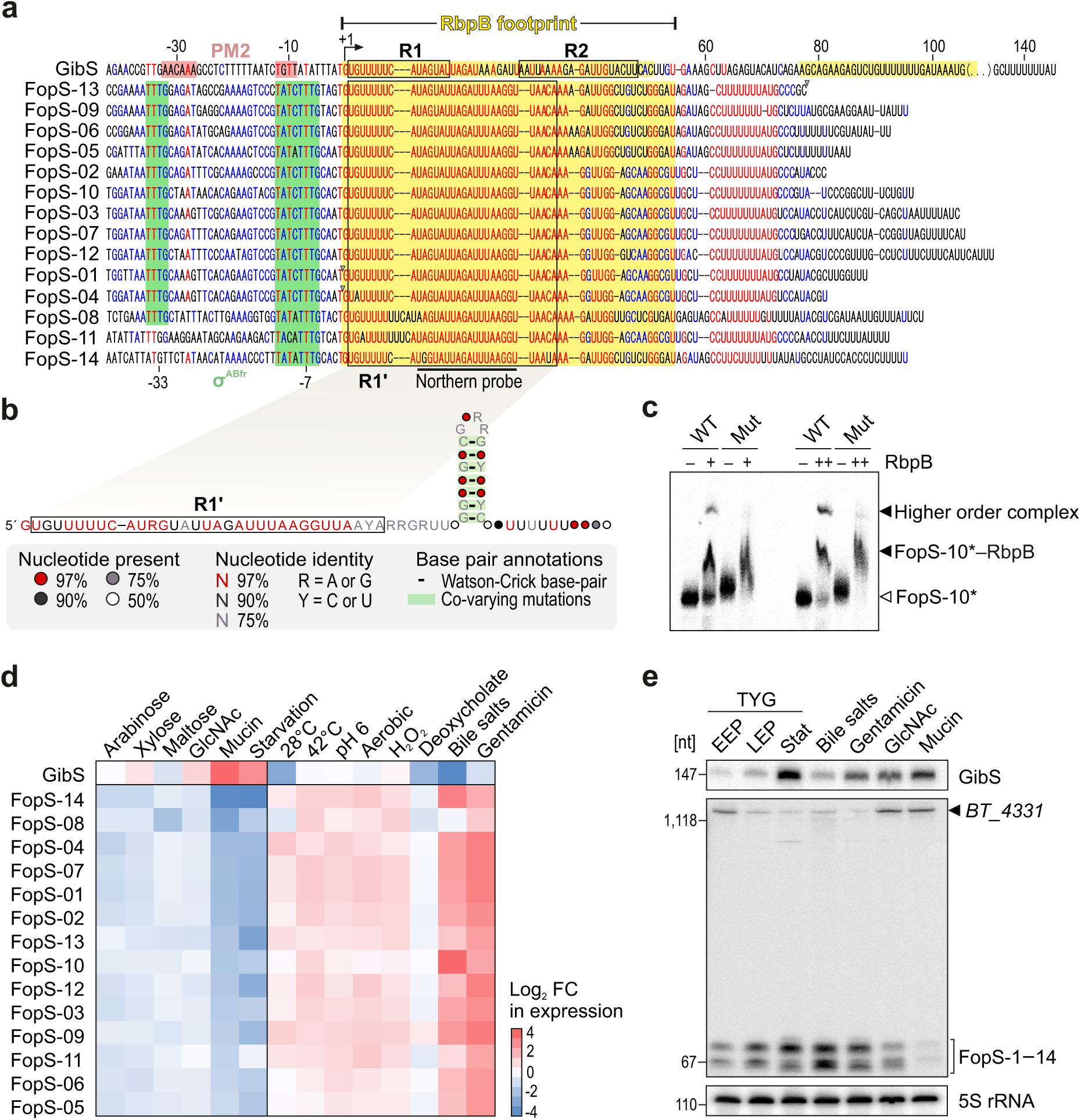
A fourteen-member family of paralogous sRNAs associates with RbpB. **a,** Sequence alignment of the R1-containing sRNA genes from *B*. *thetaiotaomicron* VPI-5284. Red and blue letters indicate highly conserved and less-conserved ribonucleobases, respectively. The numbers denote the position relative to the 5’ end of GibS (bent arrow). Canonical σ^ABfr^ promoters ^36^ (green boxes), promoter motif 2 ^11^ (‘PM2’; pale red boxes), and the R1 and R2 seeds (as determined in ^11^) are indicated. R1’ refers to the 3’-extended seed of FopS sRNAs (see Fig. 4d). The originally annotated 5’ ends of FopS-01 (BTnc025) and FopS-04 (BTnc032), as well as the 3’ end of FopS-13 (BTnc188) were supported by only few RNA-seq reads ^11^. We therefore manually curated the transcript boundaries to the previously identified secondary transcription start sites ^11^ in case of FopS-01 and FopS-04, and a 3’ end downstream of a predicted terminator hairpin in case of FopS-13 (curated sRNA boundaries marked by grey triangles). **b,** FopS consensus structure predicted using the Webserver for Aligning structural RNAs (WAR) ^32^. Secondary structure was visualized using the R2R software ^83^ and deposited in the Rfam database ^84^ (RFID; accession number pending). The R1’ sequence is boxed. **c,** *In vitro* - transcribed, radioactively labeled FopS-10 was incubated for 1 hour with two defined concentrations of recombinant RbpB (‘+’: 10 µM; ‘++’: 20 µM) and the resulting complexes resolved on denaturing gels. ‘WT’ refers to the wild-type sequence of this sRNA and ‘Mut’ to a FopS-10 variant with an inverted RbpB-target sequence (see Suppl. Fig. S6d). **d, e,** Expression profiling of GibS and the FopS sRNAs. Relative expression of the indicated sRNAs over a range of various carbon sources (relative to growth in minimal medium with glucose as the sole carbon source) and stress conditions (relative to an unstressed control sample) based on RNA-seq data from ^12^ (**d**) and northern blot-based validation (**e**). EEP, early exponential phase; MEP, mid-exponential phase; stat, stationary phase in TYG. The annealing site of the FopS-specific northern probe is indicated in panel a. The upper band corresponds to the primary transcript of the 3’ UTR-derived Fops-14. Positions of the marker bands are shown to the left. 5S rRNA served as loading control.

GibS was also pulled down together with RbpB (Fig. 2b). Previously, we had shown that this sRNA comprises two distinct seed regions, referred to as R1 and R2, which mediate base-pairing with—and repression of—its two direct target mRNAs ^11^. Interestingly, the FopS family members contain the R1 sequence of GibS but lack R2 (Fig. 3a). We adopted a structural RNAs alignment tool ^32^ to infer the consensus structure of the 14 FopS sRNAs of *B*. *thetaiotaomicron*. This structure comprised a single-stranded 5’ stretch followed by a Rho-independent terminator hairpin (Fig. 3b), and was deposited in the Rfam database (RF-ID pending). The conserved R1 sequence is located within the predicted single-stranded region at the 5’ end of the FopS consensus structure, as would be expected if R1 acts as a seed sequence in these sRNAs.

Electrophoretic mobility shift assay (EMSA) with recombinant *B*. *thetaiotaomicron* RbpB (Suppl. Fig. S6a) and—as a representative sRNA family member—*in*-*vitro* -transcribed FopS-10 supported the formation of stable ribonucleoprotein (RNP) particles (Fig. 3c; Suppl. Fig. S6c). The stoichiometry of the RbpB:FopS complex is likely greater than 1, as evidenced by the observed upshift and supershift. This is supported by size-exclusion chromatography analysis, in which RbpB behaved as a homodimer *in vitro* (Suppl. Fig. S6b). Moreover, and despite RbpB bound diverse sequence motifs *in vivo* (Suppl. Fig. S4a), its interaction with FopS was sequence-specific. That is, inverting the sequence of the 55 nt-long RbpB binding site (as deduced from CLIP-seq) predicted to largely maintain the secondary structure of FopS-10 (Suppl. Fig. S6d), was sufficient to abrogate formation of the higher order RNP complex (‘Mut’ in Fig. 3c; Suppl. Fig. S6c). EMSAs also validated that RbpB binds to GibS, again forming higher order complexes (Suppl. Fig. S4b, Suppl. Fig. S6e). In the case of both tested sRNAs, the affinity of the protein to its ligand was in the low micromolar range, and thus an order of magnitude lower than what is typically observed in interactions of pseudomonadotal Hfq and ProQ with their cognate sRNA partners ^33–35^. Taken altogether, these data support the CLIP-seq results and confirm that RbpB binds to FopS and GibS sRNAs in a sequence-specific and concentration-dependent manner. Our findings also suggest that RbpB forms multimers, most likely homodimers, on its sRNA ligands.

### FopS expression responds to bile salts

Oftentimes, expression profiling of a given sRNA provides a glimpse into its function. While GibS transcription is driven from a non-canonical promoter associated with stationary phase-induced genes ^11^, inspection of the regions upstream of the *fopS* genes revealed the presence of the canonical σ^ABfr^ promoter ^36^ (red or green boxes, respectively, in Fig. 3a). In accordance with an independent transcriptional activation, published RNA-seq data ^12^ showed an anticorrelated expression pattern of GibS and the FopS sRNAs (Fig. 3d). As confirmed by northern blotting (Fig. 3e), the steady-state level of GibS increased during growth in rich TYG medium up to stationary phase, but was highest when bacteria fed on GlcNAc-containing carbohydrates (as previously reported ^11^), particularly when mucin was the sole carbon source. In contrast, GibS levels dropped when bacteria were exposed to bile salts. Under those conditions, *B*. *thetaiotaomicron* induced expression of the FopS sRNAs, reflected by two, relatively distinct bands that matched the predicted length of the FopS sRNAs (two clusters of ∼65 and ∼70 nt) on a northern blot (Fig. 3e) using a probe against the universal FopS 5’ region (Fig. 3a). An additional, high molecular band of ∼1,400 nt was most likely derived from *BT_4331*, i.e. the parental transcript of the 3’ end-derived FopS-14 (Suppl. Fig. S7a).

Concentration range experiments coupled to northern blotting revealed FopS expression to peak between 0.03 and 0.1 mg/mL of bile salts (Suppl. Fig. S7b), falling within the range of their *in*-*vivo* concentrations in the human large intestine ^37,38^. The bile salt cocktail we used consisted of cholic acid (a primary bile salt) and deoxycholate (a conjugated, secondary bile salt) in equimolar ratio. However, interrogation of previously generated RNA-seq data ^12^ indicated deoxycholate alone to be insufficient to induce FopS expression (Fig. 3d). Using northern blotting, we observed that the primary bile acid alone did not strongly activate FopS expression either (Suppl. Fig. S7c). Our analysis thus further refined the FopS-inducing stimulus to physiological concentrations of bile salt mixtures composed of both, primary and secondary bile acids. Responsiveness to this *in*-*vivo*-relevant stimulus corroborates the relevance of the identified sRNA-RBP cluster for *B*. *thetaiotaomicron* within its host niche.

### Established GibS targets are refractory to FopS

We next sought to functionally characterize the FopS sRNA family. Previously we found that GibS represses *BT_0771* and *BT_3893*, which code for a glucan-branching enzyme that belongs to the glycoside hydrolase-13 family (http://www.cazy.org/) and a hypothetical protein, respectively ^11^. Mechanistically, GibS binds the translation initiation region of *BT_0771* mRNA and of *BT_3893* mRNA, the latter of which by engaging both the R1 and R2 seeds ^11^. To assess if these GibS targets are also regulated by the FopS cluster, we compared their steady-state transcript levels between wild-type and three independent, randomly chosen *fopS* deletion mutants (Δ*fopS-09*, Δ*fopS-10*, Δ*fopS-13*). In stationary phase TYG cultures, endogenous GibS and FopS sRNAs were relatively highly expressed (Fig. 3e). However, both established GibS target mRNAs were derepressed only in the absence of GibS and not in the absence of individual FopS sRNAs (Fig. 4a). Consistent with the notion that GibS is the primary regulator of these mRNAs, *in*-*vitro* -transcribed 5’ ends of *BT_3893* and *BT_0771* mRNAs annealed with radiolabeled GibS, but not (*BT_3893*) or less efficiently (*BT_0771*) with FopS-10 (Suppl. Fig. S8a). Therefore, although functional redundancy amongst the paralogous sRNAs might partially compensate for effects derived from single *fopS* deletion, it appears from EMSAs that FopS sRNAs are relatively ineffective in regulating GibS targets, despite possessing one of the two seed regions of GibS.

**Figure 4:**
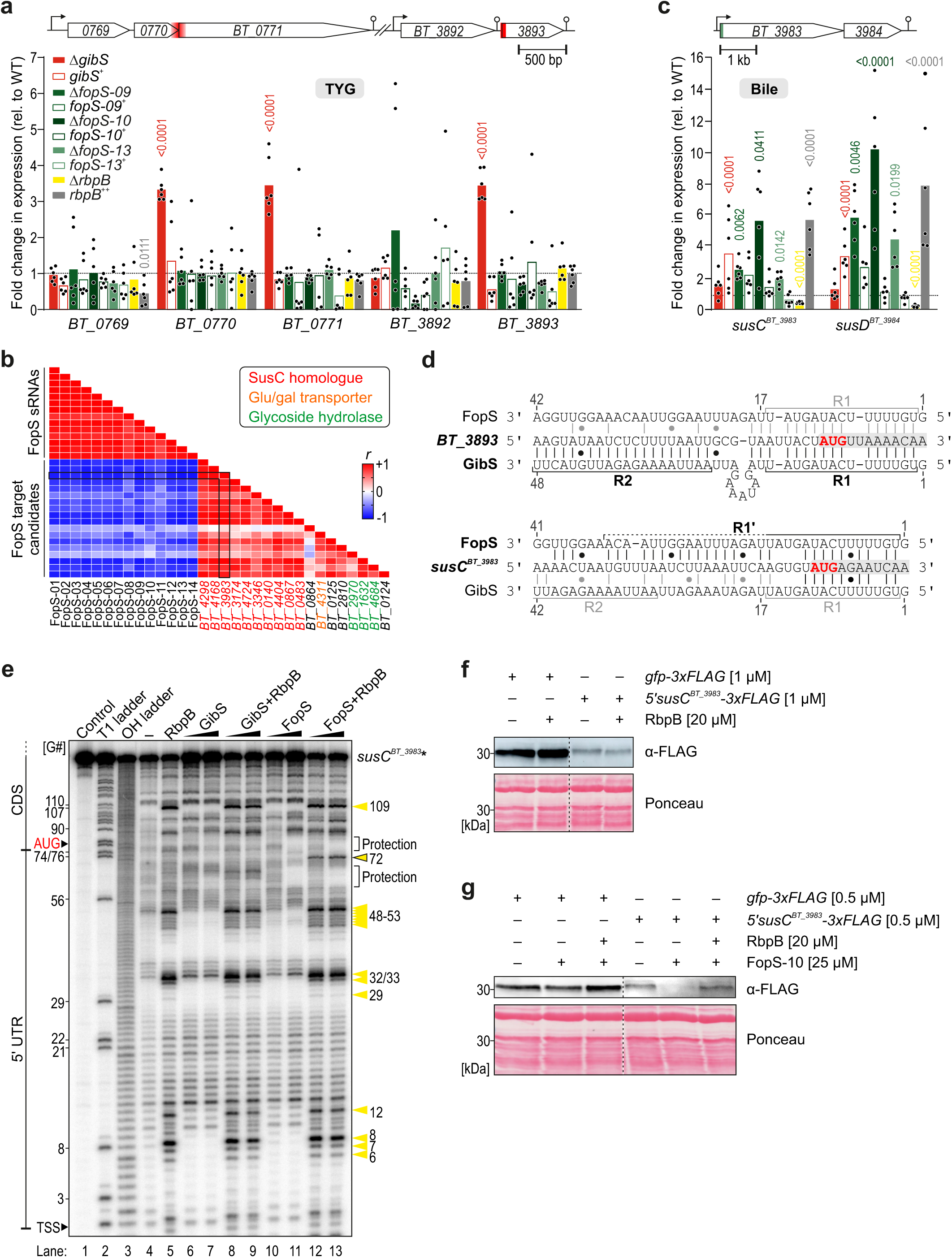
FopS sRNAs and RbpB constitute a post-transcriptional layer of PUL control. **a,** qRT-PCR-based profiling of the two direct targets of GibS, *BT_0771* and *BT_3893*, in the indicated strains grown to stationary phase in TYG ^11^. As reported previously, GibS-mediated control of the *BT_0771* -harboring polycistronic RNA extended to the adjacent *BT_0770* ^11^. Normalization was against 16S rRNA. Bars denote the mean from six replicate measurements (represented as single dots). Significance was assessed using two-way ANOVA (Sidak’s multiple comparisons test; *p* values for significant [*p* < 0.05] expression changes relative to the isogenic wild-type [dashed horizontal line] are given). Target locus representations are given at the top, with the experimentally mapped GibS binding site within *BT_3893* ^11^ (see also panel **d**) and the 156 nt-region within *BT_0771* shown previously to interact with GibS in a gel retardation assay ^11^ marked in red. **b,** Anticorrelation between the steady-state mRNA levels of *in*-*silico* -predicted FopS targets and individual FopS sRNAs upon growth on different nutrient sources and under defined stress conditions based on RNA-seq data from ^12^. Plotted in the chart are the Pearson’s correlation coefficients (*r*) for the individual sRNA-mRNA pairs. The order of genes on the y-axis is the same as on the x-axis. As an example, the (anti-)correlation of *BT_3983* with all FopS family members and other predicted target genes is boxed. **c,** qRT-PCR measurement of the steady-state mRNA levels of the predicted FopS target operon *susCD^BT_3983—84^*, analogous to panel **a**, except that strains were exposed for 2 h to 0.5 mg/mL of bile salts prior to RNA extraction and analysis. Target locus representation at the top, with FopS binding site (determined in panels **d, e**) in green. **d,** RNA:RNA interaction predictions for the GibS and FopS sRNAs (FopS-10 is displayed as a representative FopS member) with their respective targets. The start codon is labeled in red and the gray shading denotes the coding region. R1, R2: seed regions of GibS as defined in ^11^; R1’: 3’-extended seed region of the FopS’s compared to GibS R1. Coordinates are relative to the sRNAs’ 5’ ends. **e,** In-line probing of 0.2 pmol ^32^P-labeled *susC^BT_3983^* (region –69 to +117 relative to the start codon) in the absence or presence of either 20 nM or 200 nM GibS, FopS-10, or both sRNAs, and 20 µM of recombinant RbpB. ‘Control’ and ‘–‘ refer to the radiolabeled RNA substrate in water or in-line buffer, respectively, and partially RNase T1-(‘T1’) or alkali-digested (‘OH’) substrate were included as ladders. **f–g,** *In* -*vitro* translation assay of T7-transcribed fusion mRNAs *5’ susC ^BT_3983^-3xFLAG* (native *B*. *thetaiotaomicron* RBS) or *gfp-3xFLAG* (negative control; *E*. *coli* optimized RBS). 1 µM (**f**) or 0.5 µM (**g**) of fusion mRNA were *in vitro* -translated with reconstituted *E*. *coli* 70S ribosomes. Translation was performed in the absence or presence of 20 µM purified RbpB (**f**, **g**) and of 25 µM *in vitro* -transcribed FopS-10 sRNA (**g**). Translation products were resolved and detected via western blotting. Ponceau S staining of the proteinaceous translation reagents served as loading control.

### FopS sRNAs repress PUL72 by base-pairing to the cognate susCD operon

To identify the *bona fide* function of the FopS, we applied the IntaRNA algorithm ^39^ using the consensus FopS 5’ sequence (position 1 to 21) as query (Suppl. Fig. S8b). The proposed target candidates were strongly enriched in mRNAs encoding outer membrane proteins, particularly SusC-like transporters of PUL systems (green arrowheads in Suppl. Fig. S8b). Interrogation of available RNA-seq data ^12^ revealed a strong anticorrelation between the expression of individual FopS sRNAs and that of their *in*-*silico* -predicted, common targets (Fig. 4b), implying negative regulation to prevail amongst putative FopS-mediated activities.

For further characterization of FopS-mediated target control, we henceforth focused on PUL72, which is functionally and structurally characterized ^19,40,41^ and for which the glycan substrate—namely high mannose *N*-glycan—is known ^40^. IntaRNA predictions were corroborated by the steady-state transcript levels of *BT_3983*, encoding the SusC homologue of this PUL. In line with a repressive effect of the FopS sRNAs on this SusC-like transporter, its mRNA level was elevated in individual Δ*fopS* strains during growth under bile stress, and returned to basal level when *fopS* was complemented in *trans* (Fig. 4c). The mRNA of the PUL72-encoded surface glycan-binding SusD homologue (*BT_3984* ; encoded downstream in the same polycistronic mRNA as the SusC homologue; Fig. 4c) showed a similar FopS-dependent expression profile, implicating that FopS binding to the 5’ region of *susC^BT_3983^* represses both genes in this operon. Deletion of *gibS* did not affect the steady-state levels of any of these transcripts.

*In* -*silico* prediction of the RNA-RNA interaction suggested that the FopS 5’ end base-paired with the translation initiation region of *susC^BT_3983^* (Fig. 4d). This involves a 3’-extended version of the R1 region within FopS (that we term R1’)—a sequence that includes several mismatches in GibS (Fig. 3a), which might explain the divergent outcomes of deleting these sRNAs on levels of the *susC^BT_3983^* mRNA (Fig. 4c). Indeed, inline probing of the radiolabeled 5’ region of the *susC^BT_3983^* mRNA (between the 5’ end and the 69^th^ codon of the coding sequence) upon incubation with increasing concentrations of *in*-*vitro* -transcribed GibS or FopS-10 showed the predicted targeting site is shielded by FopS-10, but not by GibS (Fig. 4e; compare lanes 10 and 11, or 6 and 7, respectively, with lane 4). EMSA likewise confirmed binding of FopS-10 to the 5’ region of *susC^BT_3983^* mRNA, whereas the addition of up to 1 µM of GibS did not result in an upshift of the radioactively labeled mRNA fragment (Suppl. Fig. S8c). Taken together, these data suggest that the FopS sRNAs—but not GibS—repress PUL72 and possibly additional PUL systems (Fig. 4b, Suppl. Fig. S8b) by direct base-pairing to the 5’ region of the mRNA of the corresponding *susC* homologue.

### FopS-10 suppresses translation initiation of susC ^BT_3983^ and is antagonized by RbpB

Current paradigm holds that RNA binding proteins can repress target mRNA translation by facilitating sRNA binding, in which the specificity of the binding is determined by the base-pairing between the sRNA and the target. To investigate whether RbpB assumes similar roles, we performed three-component EMSAs. Compared to the affinities of GibS and FopS-10 to their respective targets in the absence of RbpB (black curves in Suppl. Fig. S9a), supplementation of a fixed concentration (1 µM) of the protein did not affect sRNA-mRNA duplex formation efficiency *in vitro* (yellow curves in Suppl. Fig. S9a). Gradually increasing the RbpB concentration to up to 20 µM suggested the formation of a trimeric complex consisting of GibS, its target mRNA (*BT_3893*), and RbpB, whereas the protein was not engaged in a stable complex with FopS-10 and its mRNA target (*susC^BT_3983^*) under the same conditions (Suppl. Fig. S9b, c). Unexpectedly, high concentrations of the protein (>5 µM) titrated FopS-10 away from its target *in vitro* (Suppl. Fig. S9c). Likewise, overexpression of the protein *in vivo* resulted in increased, and *rbpB* depletion in reduced steady-state levels of the *susCD* operon of PUL72 (Fig. 4c), suggesting that RbpB counteracts FopS-mediated target repression.

Moreover, EMSAs also revealed that the *susC^BT_3983^* and *BT_3893* mRNAs themselves are ligands of RbpB—even in the respective absence of FopS or GibS (Suppl. Fig. S9b, c). Note that none of these genes was expressed during mid-exponential growth in rich medium ^11^, providing a plausible explanation why the corresponding mRNAs were not recovered in the CLIP-seq experiment (Fig. 2). This sRNA-independent interaction was further supported by inline probing, revealing substantial structural rearrangements in the 5’ UTR in the presence of RbpB (compare lanes 4 and 5 in Fig. 4e). Inline probing of the interaction between RbpB and the 5’ region of the *susC^BT_3983^* mRNA in the additional presence of FopS-10 uncovered one further structural change (a band present only in lanes 12 and 13 in Fig. 4e). This rearrangement occurred at position 72 relative to the 5’ end, which coincides with position –6 with respect to the start codon of the mRNA (see Suppl. Fig. S10 for a model of the structural rearrangements) and thus, falls within the critical window for efficient translation initiation in the *Bacteroidota* ^42^.

*In* -*vitro* translation of a *susC^BT_3983^-3xFLAG* mRNA template was unaffected by the addition of recombinant RbpB alone (Fig. 4f). In contrast, addition of FopS-10 to the reaction mix abrogated target protein synthesis (Fig. 4g). In the additional presence of recombinant RbpB, FopS-10-mediated translational repression was relieved (Fig. 4g). Translation of a control template, consisting of an *Escherichia coli* ribosome-binding site fused to the 5’ region of the GFP open reading frame and a triple-FLAG tag, was largely unchanged in analogously prepared reactions (Fig. 4f, g). Based on these data, we propose that FopS-10 sRNA blocks translation of the *susC^BT_3983^* mRNA by binding adjacent to the start codon and providing steric hindrance to the initiating ribosome. This repression is counteracted by RbpB, possibly by titrating FopS-10 away from its target mRNA and/or by opening up the translational enhancer region of *susC^BT_3983^* that is otherwise occluded by FopS-10 (Suppl. Fig. S10). Altogether, our mechanistic data suggest that, in contrast to well-characterized RNA binding proteins, RbpB assumes dual roles in facilitating and excluding the binding of sRNA to a wide range of target transcripts, thus serving as a central hub in *Bacteroides* metabolic control.

## DISCUSSION

The composition of the gut microbial community and, therefore, the pivotal functions these microbes provide to human health hinges on their ability to persist in the face of nutritional fluctuation. Understanding how gut commensals adapt their metabolism to the daily variations in feeding rhythm and types of diet has thus become an important branch of microbiota research ^43,44^. The predominant bacterial genus in the healthy human gut microbiota, *Bacteroides*, possesses dozens of multi-protein complexes encoded on dedicated genomic loci— the PULs—to bind, clip, and import specific polysaccharides. Historically, PUL function was first studied in *B*. *thetaiotaomicron* ^45,46^, serving as a paradigm for polysaccharide breakdown in the human microbiome, with practical applications ^47^. Complementing previous studies that focused on transcriptional control mechanisms ^7–10,20,48,49^, the present work revealed a remarkably complex RNA-based regulatory circuit governing PUL regulation in *B*. *thetaiotaomicron*. At the heart of this network are the conserved RRM-1 protein RbpB and a family of paralogous sRNAs. Together, they constitute an elaborate post-transcriptional network to optimize fitness by orchestrating mutually exclusive expression of opposing catabolic processes (Fig. 5). Given the conservation of RRM-1 proteins ^15^ and R1 sequence-containing sRNAs (Suppl. Fig. S5a) across the *Bacteroidota* phylum, this regulatory circuit is likely prevalent in a substantial fraction of mammalian intestinal microbiota members.

**Figure 5:**
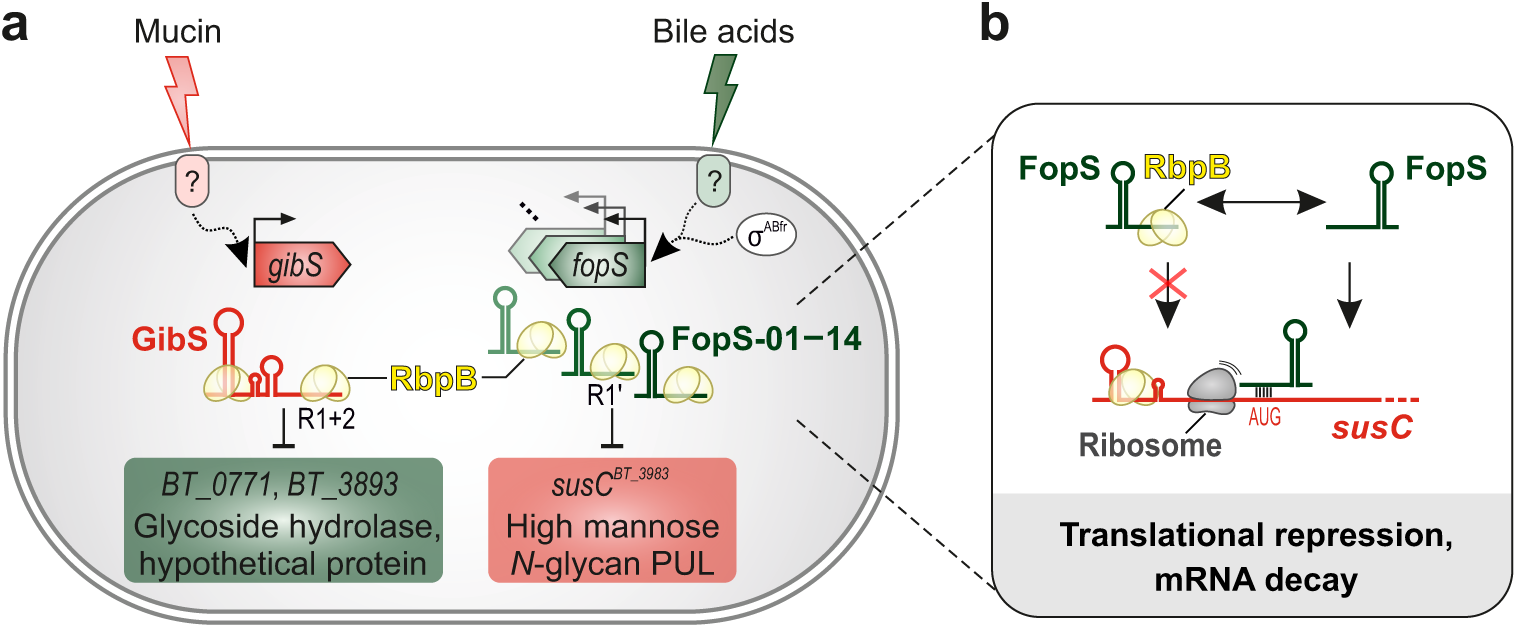
Working model for the functional diversification between RbpB-associated sRNAs—GibS and the FopS sRNA cluster—in *B*. *thetaiotaomicron* metabolism control. **a,** GibS transcription is activated during growth on GlcNAc-containing glycans (especially mucin) through an elusive transcriptional factor recognizing a previously identified promoter motif in the *gibS* promoter (‘PM2’ ^11^). GibS associates with RbpB (two distinct CLIP peaks) and represses the glycoside hydrolase 13 (GH13)-containing protein BT_0771 as well as the uncharacterized protein BT_3893 through base-pairing with its two distinct seeds (R1 and R2) to the translation initiation region of the corresponding mRNAs. In contrast, transcription of the 14 FopS sRNAs is driven from σ^ABfr^-dependent promoters and likely activated by an unknown transcriptional regulator during exposure to bile salts and potentially other membrane-assaulting stresses. The FopS sRNAs also associate with RbpB (each one CLIP peak in the 5’ portion of the sRNAs). Through their extended seed sequence R1’, the FopS sRNAs bind and repress translation of the mRNA for the outer membrane porin SusC^BT_3983^ of PUL72 and potentially additional SusCD homologs. **b,** The predicted role of RbpB in this context. *In vitro*, FopS-10 inhibited SusC^BT-3983^ translation and addition of recombinant RbpB was sufficient to overcome repression. *In vivo*, expression of *susC^BT_3983^* was upregulated in an *rbpB* overexpression strain and decreased in the absence of the protein. This implies that RbpB counteracts FopS-mediated target repression.

Paralogous sRNAs are ubiquitously present in the bacterial kingdom, but have mostly been studied in *Pseudomonadota* (formerly *Proteobacteria*) and *Bacillota* (*Firmicutes*) ^50^. In these phyla, multicopy sRNAs are frequently involved in regulating lifestyle transitions. *E*. *coli* and *Salmonella enterica*, for example, encode the paralogous sRNA pair OmrA/B to reinforce the transition between motility and biofilm formation ^51–54^. A pair of sibling sRNAs governs the switch from cataplerotic to anaplerotic metabolism in *Neisseria* ^55–58^. The Qrr sRNA family integrates quorum-sensing signals to coordinate biofilm formation and virulence in *Vibrio* spp. ^59,60^. The five csRNAs of *Streptococcus pneumonia* repress competence ^61,62^ and no less than seven sRNA paralogs (LhrC1-7) regulate virulence programs in the food-borne pathogen *Listeria monocytogenes* ^63,64^.

What are the functional benefits that prompt bacteria to maintain multicopy sRNAs in their genome despite the evolutionary pressure to minimize energy expenditure? Since they share a substantial degree of sequence and structure, sRNA paralogs typically regulate an overlapping set of target genes, yet may also have exclusive targets. Non-redundant regulatory functions may additionally arise from differential sRNA activation, e.g. the expression of paralogous sRNAs may be governed by different σ factors. Within the FopS sRNA cluster (excluding GibS), we did not observe strong indication of sub-diversification among individual family members, except for a few SNPs in the R1’ seed region of four FopS sRNAs (FopS-04, FopS-08, FopS-11, FopS-14; Fig. 3a). However, coexpression of the FopS sRNAs might have implications for dosage-dependent effects. For example, multiplexing FopS transcription across multiple loci could minimize the response time upon sensing a certain stress or nutrient cue, and rapidly rearrange cell-surface components needed to cope with hostile conditions and/or outpace metabolic competitors. Of note, the FopS cluster is not the only multicopy sRNA family in *Bacteroides* that is associated with RbpB. For example, the second enriched sequence motif within its ligands (Suppl. Fig. S4a) is contained in another paralogous sRNA family, consisting of six members that are each encoded adjacent to transposases/invertases. CRISPR-based combinatorial perturbation screens that build upon existing technology ^65^ might in the future be leveraged to dissect additive effects and functional redundancies within the FopS cluster and other paralogous sRNA families in *Bacteroides* and beyond.

The *Bacteroidota* diverged from the common line of eubacterial descent before other groups ^66^. Consequently, these bacteria are fundamentally different from the other major Gram-negative phylum, the *Pseudomonadota*, with its long-standing model species *E*. *coli* and *S*. *enterica*. For example, *Bacteroides* spp. evolved unique transcription ^67^ and translation ^68^ initiation signals. Importantly, although they encode hundreds of noncoding RNAs ^11–14^, these bacteria lack the classical chaperones of pseudomonadotal sRNAs ^69^. Despite recent attempts to infer *Bacteroidota* RNA chaperones from *in*-*silico* prediction and *in*-*vitro* experimentation ^14,15^, up to now the only global RBP that has been functionally characterized in this phylum remained the transcription termination factor Rho ^70^.

Applying CLIP-seq, we demonstrated that *B*. *thetaiotaomicron* RbpB directly associates with >200 mRNAs and >60 noncoding RNAs. Among the latter are the functionally characterized sRNAs GibS ^11^, MasB ^12^, and BatR ^65^, which all act via base-pairing to complementary stretches near the start codon of their respective target mRNAs. This warrants future investigation into the role of RbpB in these regulations, which may reveal new and potentially generalizable sRNA-mediated regulatory principles. Besides, we observed sRNA-independent binding of RbpB to the 5’ UTRs of selected mRNAs *in vitro* and according to our CLIP-seq data, RbpB preferentially binds to AU-rich sequences that are frequently found around the *Bacteroides* ribosome-binding site ^68^. A pathway enrichment analysis further indicated translation-associated processes to be overrepresented amongst the functional annotations of RbpB mRNA ligands. In the one tested example, the addition of recombinant RbpB did not alter the translational output of *susC^BT_3983^* in the absence of a repressing sRNA. As *Bacteroides* RNA elements are increasingly recognized as targets for microbiome editing^71^, a profound knowledge of the mode of action and the molecular players involved in endogenous RNA-mediated processes in these bacteria is urgently needed.

In summary, the present work uncovered a large RNA network controlling PUL translation in *Bacteroides* —genetic loci that are important for efficient colonization of the mammalian gut and previously known to be regulated transcriptionally. More generally, our study serves as a model for RNA-mediated metabolism control by gut commensals and proposes RRM-1 domain-containing proteins as excellent candidates to identify additional post-transcriptional hubs in the microbiota. This knowledge could be key to exploiting this microbial consortium for therapy against infectious diseases and intestinal disorders.

## EXPERIMENTAL PROCEDURES

### Bacterial cultivation and genetics

Liquid cultures of *Bacteroides thetaiotaomicron* strains were prepared in complex tryptone-yeast extract-glucose (TYG) medium (20⍰g⍰L-1 tryptone, 10⍰g⍰L-1 yeast extract, 0.5% glucose, 5⍰mg⍰L-1 hemin, 1⍰g⍰L-1 cysteine, 0.0008% CaCl_2_, 19.2⍰mg⍰L-1 MgSO_4_·7H_2_O, 40⍰mg⍰L-1 KH_2_PO_4_, 40⍰mg⍰L-1 K_2_HPO_4_, 80⍰mg⍰L-1 NaCl, 0.2% NaHCO_3_) or minimal medium (1⍰g⍰L-1 L-cysteine, 5⍰mg⍰L-1 hemin, 20⍰mg⍰L-1 L-methionine, 4.17⍰mg⍰L-1 FeSO_4_, 0.2% NaHCO_3_, 0.9⍰g⍰L-1 KH_2_PO_4_, 0.02⍰g⍰L-1 MgCl_2_·6H_2_O, 0.026⍰g⍰L-1 CaCl_2_·2H_2_O, 0.001⍰g⍰L-1 CoCl_2_·6H_2_O, 0.01⍰g⍰L-1 MnCl_2_·4H_2_O, 0.5⍰g⍰L-1 NH_4_Cl, 0.25⍰g⍰L-1 Na_2_SO_4_) supplemented with 0.5% of the indicated carbon sources. For propagation on plates, brain heart infusion-supplemented (BHIS) agar (52⍰g⍰L-1 BHI agar powder, 1⍰g⍰L-1 cysteine, 5⍰mg⍰L-1 hemin, 0.2% NaHCO_3_) was used. Cultures were incubated at 37°C in an anaerobic chamber (Coy Laboratory Products) in presence of an anoxic gas mix of 85% N_2_, 10% CO_2_, 5% H_2_. *Escherichia coli* strains were cultured aerobically in Luria-Bertani (LB) broth (10 g L-1 tryptone, 5 gL-1 yeast extract, 10 g L-1 NaCl) at 37°C with shaking or statically on LB agar plates. All strains, plasmids, and oligonucleotides used in this study are listed in Supplementary Table 2.

### Mouse experiments

C57BL/6J wild-type (cat# 000664), were obtained from Jackson Laboratory. Mice were housed in sterile cages under specific pathogen-free conditions on a 12-hour light cycle, with *ad libitum* access to food and sterile water at Vanderbilt University Medical Center. Seven to nine-week-old male mice were randomly assigned into treatment groups before the experiment. One week before antibiotic treatment, mice were switched to a fiber-free diet (TD.130343) or remained on a control diet 5010 (LabDiet 0001326) until the end of the experiment. Antibiotic cocktails (ampicillin [Sigma-Aldrich], metronidazole [Sigma-Aldrich], vancomycin [Chem Impex International], and neomycin [Sigma-Aldrich]; 5 mg of each per mouse) were administered by oral gavage daily for 5 days. After antibiotic treatment, mice were inoculated with an equal mixture of 0.5 × 10^9^ CFU of the *B*. *thetaiotaomicron* wild-type strain and 0.5 × 10^9^ CFU of indicated mutants for 6 days. After euthanasia, cecal and colonic tissue was collected in sterile PBS, and the abundance of *B*. *thetaiotaomicron* strains was quantified by plating serial-diluted intestinal contents on selective agar.

### UV cross-linking and immunoprecipitation (CLIP)

CLIP-experiments were performed as described in ^72^ with minor modifications. In brief, bacterial cultures were grown to mid-exponential phase (OD_600_ = 2.0) in TYG. In total, 200 OD equivalents per condition (cross-linked, non-cross-linked) were harvested and irradiated with UV light (254 nm) at 800 mJ/cm^2^ on a 22 cm x 22 cm plastic tray. Cells were pelleted (4,000 g, 40 min, 4°C) and snap-frozen in liquid nitrogen. Each pellet was resuspended in 800 µL of NP-T buffer (50 mM NaH_2_PO_4_, 300 mM NaCl, 0.05% Tween 20, adjusted to pH 8.0). Cells were lysed mechanically in a standard Retsch apparatus (30 1/s, 10 min), using 1 mL of 0.1 mm glass beads for grinding. To clear lysates from beads, samples were centrifuged twice at 16,000 x g for each 15 min at 4°C.

Cleared lysates were mixed with equal volumes of NP-T buffer containing 8 M urea and incubated for 5 min at 65°C, with shaking at 900 rpm. Each lysate was then diluted (1:10) in pre-cooled NP-T buffer. Subsequently, 30 µL of anti-Flag M2 magnetic beads (Sigma-Aldrich) were washed and equilibrated in 800 µL of NP-T buffer, added to the samples, and incubated for 1 h at 4°C, rotating. Beads were collected by centrifugation at 1,500 g for 1 min at 4°C and washed twice with each 2 mL of high-salt buffer (50 mM NaH_2_PO_4_, 1 M NaCl, 0.05% Tween 20, adjusted to pH 8.0) and 2 mL of NP-T buffer. Each sample was resuspended in NP-T buffer, containing 1 mM MgCl_2_ and 25 U Benzonase and incubated for 10 min at 37°C, 900 rpm, followed by a 2-min incubation on ice. After washing the beads once with 1 mL of high-salt buffer and twice with 1 mL of CIP-buffer (100 mM NaCl, 50 mM Tris-HCl [pH 7.4], 10 mM MgCl_2_), 100 µL of CIP-mix containing 10 U of calf intestinal alkaline phosphatase (New England Biolabs) in CIP-buffer were added and beads incubated for 30 min at 37°C, 800 rpm. Subsequently, one wash with 500 µL of high-salt buffer and two washes with 500 µL of 1x PNK buffer (Reaction buffer A, ThermoScientific) were performed. For labeling, a PNK mix was prepared containing 98 µL of 1x PNK buffer, 10 U of T4 polynucleotide kinase (ThermoScientific), and 10 µCi γ^32^P ATP. Beads were resuspended in the PNK mix and incubated for 30 min at 37°C. Finally, 10 µL of non-radioactive ATP (1 mM) were added and samples incubated for 5 min at 37°C, before washing the beads two times with 1 mL of NP-T buffer. For elution, beads were resuspended in 15 µL of 1x protein loading buffer with 50 mM DTT and incubated for 5 min at 95°C. Magnetic beads were collected on a magnetic separator and the supernatant was transferred to a fresh tube. The elution step was repeated, the two supernatant fractions pooled and loaded (total volume of 30 µL) on a 15% SDS-polyacrylamide gel.

Protein-RNA complexes were transferred on a Portran 0.45 µm NC membrane (Amersham). The protein ladder was labeled with a radioactive marker pen and membranes were exposed to a phosphor screen overnight. The autoradiogram was visualized, printed, and aligned with the membrane to properly excise the RNA-protein complexes from the membrane. Each membrane piece was cut into smaller pieces and transferred to a low-binding tube. To each tube, 200 µL of PK mix were added, containing 2x PK buffer (100 mM Tris-HCl [pH 7.9], 10 mM EDTA, 1% SDS), 1mg/mL Proteinase K (Fermentas) and 10 U SUPERaseIN (Life Technologies), and incubated for 1 h at 37°C, 800 rpm. Additionally, 100 µL of PK buffer containing 9 M urea were added and incubated for another hour at 37°C, 800 rpm. The solution was cleared from membrane pieces and one volume of P:C:I (ROTI phenol:chloroform:isoamyl alcohol) was added. Samples were incubated in phase-lock tubes for 5 min at 30°C, 1,000 rpm, centrifuged for 12 min at 16,000 g, 4°C, and the aqueous phase transferred to a fresh tube. RNA was precipitated with three volumes of 30:1 mix (ethanol:3⍰M NaOAc, pH 6.5) and 1 µL of GlycoBlue (ThermoScientific) over night at −20°C. Subsequently, RNA was pelleted, washed with 80% (vol vol-1) ethanol, resuspended in 10 µL of H_2_O and dissolved at 65°C for 5 min, shaking (800 rpm). Samples were stored at −20°C until they were sent for sequencing.

### CLIP-seq protocol and data analysis

The preparation of cDNA libraries and high-throughput sequencing on Ilumina instrument was performed at vertis Biotechnologie AG, Freising, Germany. First, oligonucleotide adapters were ligated to the 3’ and 5’ ends, followed by a first-strand cDNA synthesis using M-MLV reverse transcriptase and the 3’ adapter as primer. The resulting cDNAs were PCR-amplified using a high-fidelity DNA polymerase with a varying number (18-24) of PCR cycles per sample. Amplified cDNA was purified using the Agencourt AMPure XP kit (Beckman Coulter Genomics). For sequencing, the samples were pooled in equimolar amounts and paired-end-sequenced on an Ilumina HiSeq system with 2×150 bp read lengths.

For analysis, raw reads were filtered and trimmed with BBDuk (paired-end mode, phred score cutoff: 20, minimal read length: 12 nt). To remove putative PCR duplicates, reads were deduplicated with FastUniq 1.1 ^73^. Mapping was done with READemption 1.0.5 pipeline using segemehl 0.3.4 ^74^. READemption align was run in paired-end mode with an accuracy of 80%. Peak calling was performed with PEAKachu pipeline version 0.2.0 (https://github.com/tbischler/PEAKachu). First, normalization factors for peak calling were calculated with an in-house *R* script as described previously in ^27^. In brief, the core positions of all libraries were isolated using an exploratory analysis of read counts summarized per position. Positions with low read counts were filtered. Normalization factors were then calculated based on the background positions which are represented by high read counts in both crosslinked and non-crosslinked libraries. PEAKachu adaptive was run in paired-end and paired-replicates mode together with the calculated normalization factors. Maximum fragment size was set to 50 and mad multiplier to 0. Only peaks with a fold-change ≥ 2 and an adjusted *p*-value ≤ 0.05 were considered significant.

As per default ^72^, we first considered only uniquely mapped reads in our CLIP-seq quantification and discarded multi-mapped reads. However, given the observed sequence similarity amongst the identified RbpB targets (the FopS sRNAs), we re-ran the analysis, this time also considering multi-mapped reads. While not affecting the overall results, this adaptation of the peak calling pipeline led to a further increase in the number of significantly enriched FopS sRNAs (from 10 FopS sRNAs when multi-mapped reads were discarded to all 14 FopS members when also these reads were included). Hence, the numbers we report herein (main text and Suppl. Table 1) are derived from the adapted (multi-mapped reads-retaining) quantification.

### Pathway enrichment analysis of CLIP-seq peaks

The genes containing at least one significant peak were checked for enrichment of gene sets belonging to a custom annotation (containing GO terms, KEGG pathways and modules, and functional information parsed from the literature ^12^). Enrichment was computed with the enricher function of the clusterProfiler R package version 3.14.3 ^75^.

### RbpB purification

Expression and purification of RbpB was performed at the recombinant protein expression facility of the Rudolf Virchow Center, Würzburg, Germany. In brief, constructs containing the coding sequence of RbpB together with a His-Sumo3 tag were cloned in an *E*. *coli* expression vector (pETM11) and transformed into *E*. *coli* Bl21 (DE3). For large-scale purification, cultures were grown to an OD_600_ of 0.6 in 8 L of LB medium, induced with 0.5 mM IPTG, and incubated over night at 18°C. Cells were resuspended in 5-10 mL of lysis buffer (150 mM NaCl, 50 mM NaH_2_PO_4_, pH 7.0, 10% sucrose, 10 mM imidazole, 1 mM TCEP, 1 mM MgCl_2_, protease inhibitor, DNase) per gram cell pellet and lysed by sonication. Lysates were cleared by centrifugation (30,000 x g, 30 min, 4°C) and incubated with 4 mL of Ni-NTA (equilibrated to lysis buffer) for 1 h at 6-8°C followed by immobilized metal affinity chromatography (IMAC) purification. Elution was performed seven times with each 4 mL of IMAC elution buffer (150 mM NaCl, 20 mM NaH_2_PO_4_, pH 7.0, 10% sucrose, 250 mM imidazole, 1 mM TCEP, 1 mM MgCl_2_).

IMAC eluates were pooled and the tag was cleaved off using SenP2 Sumo-protease. Pooled eluates were dialyzed over night against 2 L of dialysis buffer (150 mM NaCl, 20 mM NaH_2_PO_4_, pH 7.0, 10% sucrose, 1 mM TCEP, 1 mM MgCl_2_) at 6-8°C. Finally, the dialysate was concentrated (10 mL total volume), cleared by centrifugation (16,000 x g, 20 min, 4°C), and loaded on a HiLoad Superdex 75 16/600 pg for size exclusion chromatography. Eluate fractions were collected (2 mL each), pooled from two runs, and analyzed by SDS-PAGE.

### In-vitro transcription and radiolabeling of RNA

*In* -*vitro* transcription and radiolabeling of RNA was performed as described previously ^11^. Briefly, DNA templates were amplified from genomic DNA using primer pairs carrying a T7 promoter (Supplementary Table 2). The ensuing *in*-*vitro* transcription reaction was performed using the MEGAscript T7 kit (ThermoFisher Scientific). Excess DNA was removed by DNase I digestion (1 U, 37°C for 15 min) and the RNA product purified from a 6% (vol vol-1) PAA-7M urea gel using a LowRange RNA ladder (ThermoFisher Scientific) for precise sizing. The *in*-*vitro* - transcribed RNA was eluted in RNA elution buffer (0.1 M NaOAc, 0.1% SDS, 10 mM EDTA) over night on a thermoblock at 8°C and 1,400 rpm, and subsequently precipitated using ethanol:NaOAc (30:1) mix, washed with 75% ethanol and resuspended in 20 µL of water (65°C for 5 min).

For radioactive labeling, 50 pmol of the *in*-*vitro* -transcribed RNA were dephosphorylated using 25 U of calf intestine alkaline phosphatase (NEB) in a 50 µL reaction volume. After 1 h incubation at 37°C, RNA was extracted with a phenol:chloroform:isoamyl alcohol mix (P:C:I, 25:24:1) and precipitated as described above. Finally, 20 pmol of the dephosporylated RNA were 5’-end-labeled (20 µCi of ^32^P-γATP) in a 20 µL reaction for 1 h at 37°C using 1 U polynucleotide kinase (NEB). Labeled RNA was purified on a G50-column (GE Healthcare) and extracted from a PAA gel as described above.

### Electrophoretic mobility shift assay (EMSA)

Protein-RNA and RNA-RNA EMSAs were performed in a final reaction volume of each 10 µL, containing 1x RNA structure buffer (SB; Ambion), 1 µg yeast RNA (∼ 4 µM final concentration), and 5’ end-labeled RNA (4 nM final concentration). Reactions were either incubated with increasing concentrations of *in*-*vitro* -transcribed target mRNA segments (0; 8; 16; 32; 64; 128; 256; 512; 1,024 nM final concentration) or increasing concentrations of purified RbpB (0; 0.15; 0.3; 0.6; 1.25; 2.5; 5; 10; 20; 30; 40; 80 µM final concentration). Reactions were incubated for 1 h at 37°C and stopped by adding 3 µL of 5x native loading dye (0.2% bromophenol blue, 0.5x TBE, 50% glycerol) and loaded on a native 6% (vol vol-1) PAA gel in 0.5x TBE buffer at 4°C and run at 300 V for 3 h. The gel was dried at 80°C for 1 h on a Gel Dryer 583 (Bio-Rad), exposed over night, and visualized on a phosphorimager (FLA-3000 Series, Fuji).

Three-component EMSA was carried out in a 15 µL reaction volume, containing 1x RNA structure buffer (SB; Ambion), 1 µg yeast RNA (∼4 µM final concentration), and 5’ end-labeled RNA (4 nM final concentration). Depending on the experimental setup, reactions were either incubated with *in*-*vitro* -transcribed target mRNA segments or sRNAs (500 nM final concentration). Increasing concentrations of purified RbpB were added to the reactions (0; 0.15; 0.3; 0.6; 1.25; 2.5; 5; 10; 20 µM final concentrations).

### Rifampicin treatment to halt de-novo transcription

RNA stability assay was performed as described in ^14^. Briefly, single colonies of *B*. *thetaiotaomicron* strains AWS-001 (WT), AWS-323 (Δ*rbpB*), and AWS-218 (*rbpB* ^++^) were inoculated in liquid TYG medium and grown for ∼16 h, subsequently sub-cultured (1:100 dilution) and grown to mid-exponential phase (OD_600_ = 2.0). To halt *de*-*novo*-transcription, rifampicin was added (500 µg/mL final concentration). Samples were taken immediately before the addition of rifampicin (0 min) and at indicated time points after the treatment (3; 6; 9; 12; 45; 60; 90 min). Total RNA was extracted and decay assessed by northern blot analysis or quantitative real-time PCR (see below).

### RNA extraction and removal of genomic DNA

Generally, total RNA was isolated by hot phenol extraction from culture aliquots. To this end, 4 OD equivalents of culture were harvested, mixed with 20% vol. stop mix (95% vol vol−1 ethanol, 5% vol vol−1 water saturated phenol, pH >7.0), and snap-frozen in liquid nitrogen. Lysis of the bacterial cells was mediated by the addition of 600 µL of lysozyme (0.5 mg mL-1) and 60 µL of 10% SDS, followed by an incubation for 2 min at 64°C, before 66 µL of 3M NaOAc (pH 5.2) were added. For extraction, phenol was added (750 µL; Roti-Aqua phenol) and samples incubated for 6 min at 64°C, followed by the addition of 750 µL of chloroform. After centrifugation, RNA was precipitated from the aqueous phase over night at −20°C with twice the volume of 30:1 (ethanol:3⍰M NaOAc, pH 6.5). Subsequently, RNA was pelleted, washed with 75% (vol vol-1) ethanol, and resuspended in 50 µL of H_2_O. To remove contaminating genomic DNA, 40 µg of RNA were treated with 5 U of DNase I (Fermentas) and 0.5 µL of Superase-In RNase Inhibitor (Ambion) for 45-60 min at 37°C in 50 µL reaction volumes. Finally, RNA was purified with a phenol-chloroform extraction (ROTI phenol:chloroform:isoamyl alcohol) and resuspended in 30 µL of H_2_O.

### Quantitative real-time PCR

Quantitative real-time PCR was performed as described in ^11^. Briefly, a reaction mix was prepared for each well of a 96-well plate, containing 10 ng of DNase I-treated RNA, 5⍰µL of master mix (No ROX SYBR MasterMix blue dTTP kit, Takyon), 0.1⍰µL of each forward and reverse primer (10⍰µM each), and 0.08⍰µL of reverse transcriptase (One-Step Kit converter, Takyon). Analysis was on a CFX96 instrument (Biorad).

### Identification of FopS paralogs and homologs

The 14 paralogous sequences that were identified by sequence homology were manually examined for the presence of annotated transcription start and termination sites on Theta-Base^12^ and the existence of putative sORFs within each locus was excluded^76^. Originally, the annotated 5’ ends of FopS-01 (BTnc025) and FopS-04 (BTnc032), as well as the 3’ end of FopS-13 (BTnc188) were supported by only few RNA-seq reads ^11^ we therefore manually curated the transcript boundaries to the previously identified secondary transcription start sites ^11^ in case of FopS-01 and FopS-04, and a 3’ end right downstream of a predicted terminator hairpin in case of FopS-13 (curated sRNA boundaries marked by grey triangles in Fig. 3a). Based on these accurately defined transcript boundaries, we classified the individual FopS family members into intergenic sRNAs (n=9), antisense RNAs (n=4), and 3’-UTR-derived sRNAs (n=1).

### In-silico prediction of FopS sRNA consensus structure

Curation of the FopS consensus structure was done as described in detail in ^14^ and originally reported in ^77^. Briefly, alignments of the FopS sRNA sequences were generated with the WAR webserver ^32^ and the maximum consistency alignment and structure were downloaded in stockholm format. We then used the RALEE emacs “RNA editor mode” ^78^ to manually curate the alignment and optimized it with R-scape ^79^.

### Northern blotting

Northern blot analysis was carried out as previously described ^11^. In short, 5 µg total RNA were denatured for 5 min at 95°C, incubated on ice for 5 min, and separated on a 6% (vol vol-1) PAA-7M urea gel at 300 V for ∼2 h. Electroblotting of the RNA onto a Hybond-N+ membrane (GE Healthcare Life Sciences) was performed at 50 V, 4°C for 1 h. Membranes were UV-crosslinked (0.12 J cm^3^ −1), pre-hybridized in 15 mL of Hybri-Quick buffer (Carl Roth AG) at 42°C for 1 h, and incubated with ^32^P-labeled gene-specific oligonucleotides at 42°C. Blots were washed with decreasing concentrations (5×, 1×, 0.5×) of SSC buffer (20× SSC: 3⍰M NaCl, 0.3⍰M sodium citrate, pH 7.0), exposed as required, and visualized on a phosphorimager (FLA-3000 Series, Fuji).

### sRNA target prediction

To predict FopS targets, the IntaRNA program^80^ was employed. The query input consisted of a 35 nt sequence comprising the complete R1’ region at a consensus of 70% (GTGTTTTTCATAGTATTAGATTTAAGGTTAACAAA) and the program was executed using its default settings against the *B*. *thetaiotaomicron* VPI-5482 genome. Functional enrichment analysis was performed on the top 50 targets and the heat maps in Fig. S8b displays gene clusters with a DAVID^81^ enrichment score of ≥1, with colors indicating related functional categories. Genes encoding SusC-like transporters are labeled by green arrowheads. The color intensity reflects IntaRNA-derived *p*-values and the enrichment scores of the two significant clusters are given in the upper right box.

### Inline probing

In-line probing was performed as described in ^11^. Briefly, 0.2 pmol 5’ end-^32^P-labeled RNA were incubated for 40 h at room temperature in 2x in-line probing buffer (100⍰mM KCl, 20⍰mM MgCl_2_, 50⍰mM Tris-HCl, pH 8.3). Reactions were stopped by the addition of 10 µL of 2x gel-loading solution (10 M urea, 1.5 mM EDTA, pH 8.0). To prepare the RNase I ladder, 0.4 pmol 5’ end-^32^P-labeled RNA were denatured in 1x sequencing buffer (Ambion) at 95°C for 1 min. RNase TI (0.1 U) was added and incubated at 37°C for 5 min. The alkaline hydrolysis ladder was prepared by incubating 0.4 pmol 5’ end-^32^P-labeled RNA in 9 µL of 1x alkaline hydrolysis buffer (Ambion) for 5 min at 95°C. To stop the reaction, 12 µL of loading buffer II were added to both ladders and stored on ice. Samples were resolved on a 10% (vol vol-1) PAA-7 M urea sequencing gel at 45 W for 2-3 h. Gels were dried and visualized as described above.

### In-vitro translation assay and western blotting

*In* -*vitro* translation assays were performed using a reconstituted *E*. *coli* protein synthesis system (PURExpress, New England Biolabs). Reactions were performed according to manufacturer’s instruction and as previously described in ^82^ with a few modifications. In brief, 0.5 or 1 µM *in vitro* -transcribed mRNA (*5’susC^BT_3983^ -3xFLAG*, *gfp-3xFLAG*) were incubated in presence or absence with 25 or 50 µM *in vitro* -transcribed FopS-10 sRNA for 1 min at 95°C and chilled on ice for 5 min. RbpB (20 µM) was added and the reaction pre-incubated for 10 min at 37°C before the PURExpress components were added. After ∼4 h, reactions were stopped by adding 5 µL of 5x protein loading buffer and the whole sample volume (30 µL) was loaded on a 12% SDS-polyacrylamide gel. Proteins were transferred onto a Portran 0.2 µm NC membrane (Amersham) for 1.5 h, 350 mA at 4°C under semi-dry conditions. To assess equal sample loading, membranes were stained with Ponceau S (Sigma-Aldrich), visualized, and then destained with 0.1 M NaOH prior to blocking in TBS-T with 10% powdered milk (1 h, room temperature). Monoclonal anti-FLAG (Sigma-Aldrich) antibody was added (1:1,000 in TBS-T with 10% powdered milk) and incubated over night at 4°C, with shaking. Membranes were washed three times in TBS-T for each 10 min and incubated with anti-mouse IgG, HRP (ThermoFisherScientific) diluted 1:10,000 in TBS-T with 10% powdered milk for 1 h at room temperature. After a short rinsing step, ECL detection substrate (Amersham) was added and HRP activity detected using a CCD imager (Amersham ImageQuant 800 systems).

### Data availability

Sequencing data are available at NCBI Gene Expression Omnibus (http://www.ncbi.nlm.nih.gov/geo) under the accession number GSE244816.

### Code availability

Core software central to the conclusions drawn in this study are publicly available and their usage parameters described in the appropriate sections above. The CLIP-seq normalization script was previously described ^72^ and is available at github (https://github.com/lbarquist/norclip).

## SUPPLEMENTARY CAPTIONS

**Supplementary Figure S1:**
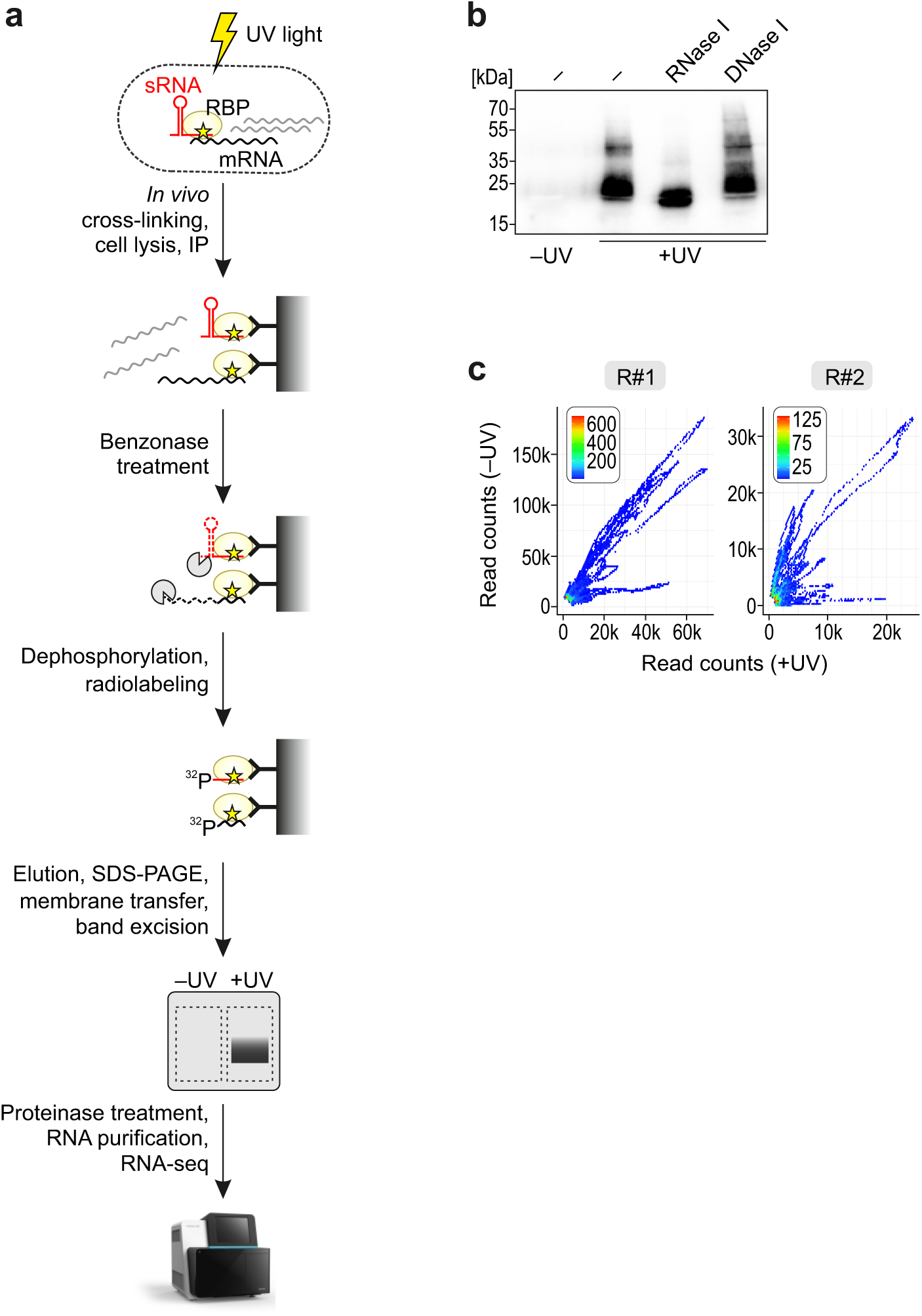
RbpB CLIP-seq. **a,** Schematic workflow of the CLIP-seq procedure to identify bacterial RBPs. Cells are irradiated with UV light (254 nm) to covalently bond protein and RNA (yellow asterisk). Upon lysis and immunoprecipitation using antibodies against the FLAG-epitope, RNP complexes are partially degraded by benzonase and subjected to radioactive labeling with polynucleotide kinase. RNPs are separated via SDS-PAGE, transferred to a nitrocellulose membrane, and the radioactive section of the membrane as well as the corresponding area in the non-cross-linked control are excised. Protein-bound RNA is released from the membrane upon proteinase K digestion and further purified through phenol:chloroform:isoamyl alcohol extraction. For RNA-seq, adapters are ligated to the RNA fragments followed by reverse transcription of RNA into cDNA, and PCR amplification. **b,** Testing sensitivity of RbpB ligands to RNase I and DNase I. Shown is the autoradiogram of a CLIP assay performed with samples treated with either RNase I or DNase I. **c,** Frequency plots of matched cross-linked and background samples for the two independently performed experiments. Plotted are the read counts per genetic feature in the cross-linked (x-axes) and non-cross-linked (y-axes) samples and the coloring refers to the frequency of each x–y pair.

**Supplementary Figure S2:**
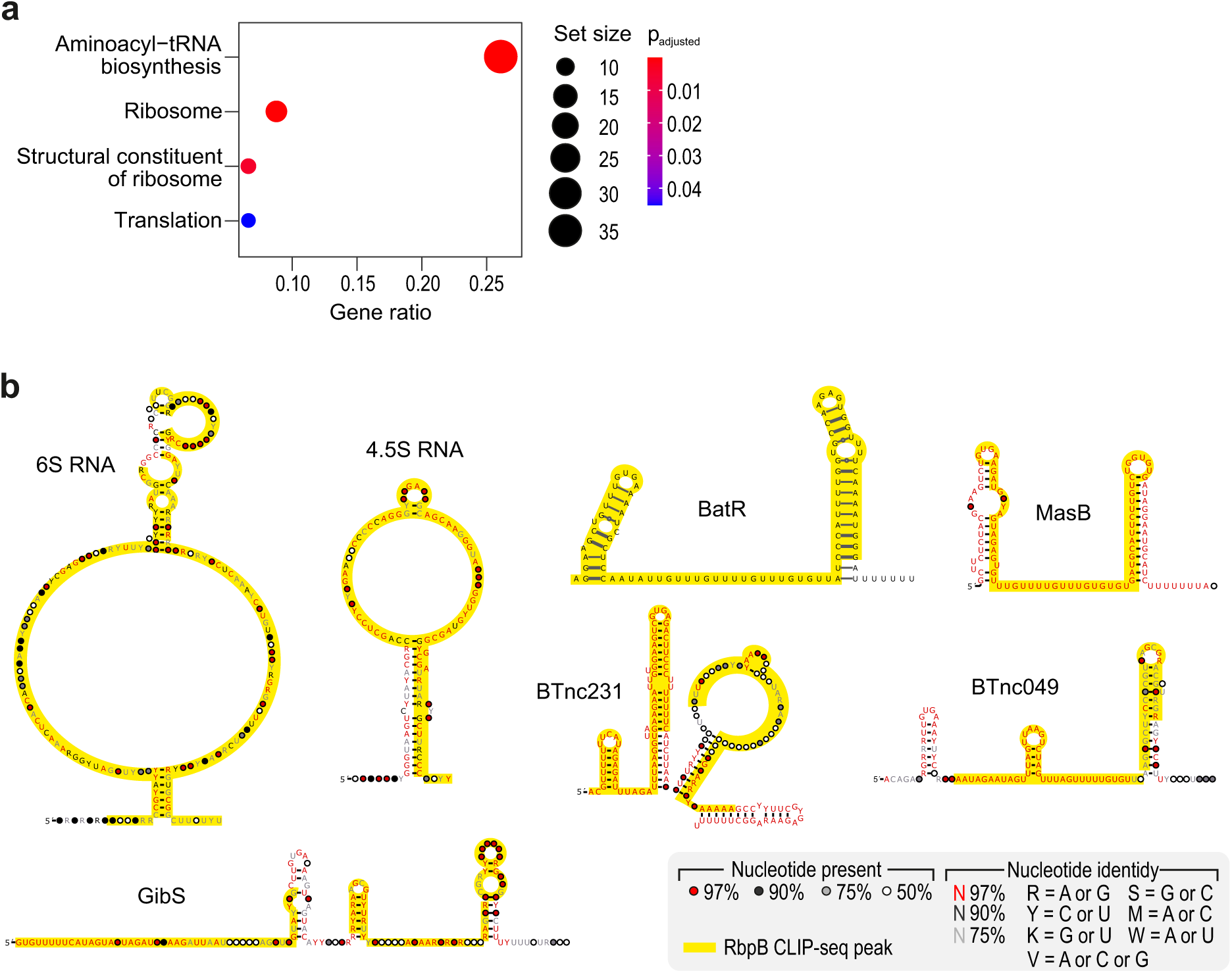
RpbB mRNA ligands are enriched for translational processesand RbpB peaks fall within single-stranded regions of sRNAs. **a,** Pathway enrichment analysis of the RbpB-bound mRNAs. Genes with at least one significant CLIP peak were checked for enrichment of gene sets belonging to a custom annotation (see Methods section). Set size indicates the number of genes in the gene set that contain a CLIP peak and gene ratio is defined as the set size divided by the total number of genes with CLIP peaks. **b,** Binding sites of RbpB within noncoding RNAs for which the secondary structure is known, namely the 4.5S and 6S housekeeping RNAs, the known *trans* -acting sRNAs GibS, MasB, BatR, and the regulatory sRNA candidates BTnc049 and BTnc231. Secondary structure information from ^14,65^.

**Supplementary Figure S3:**
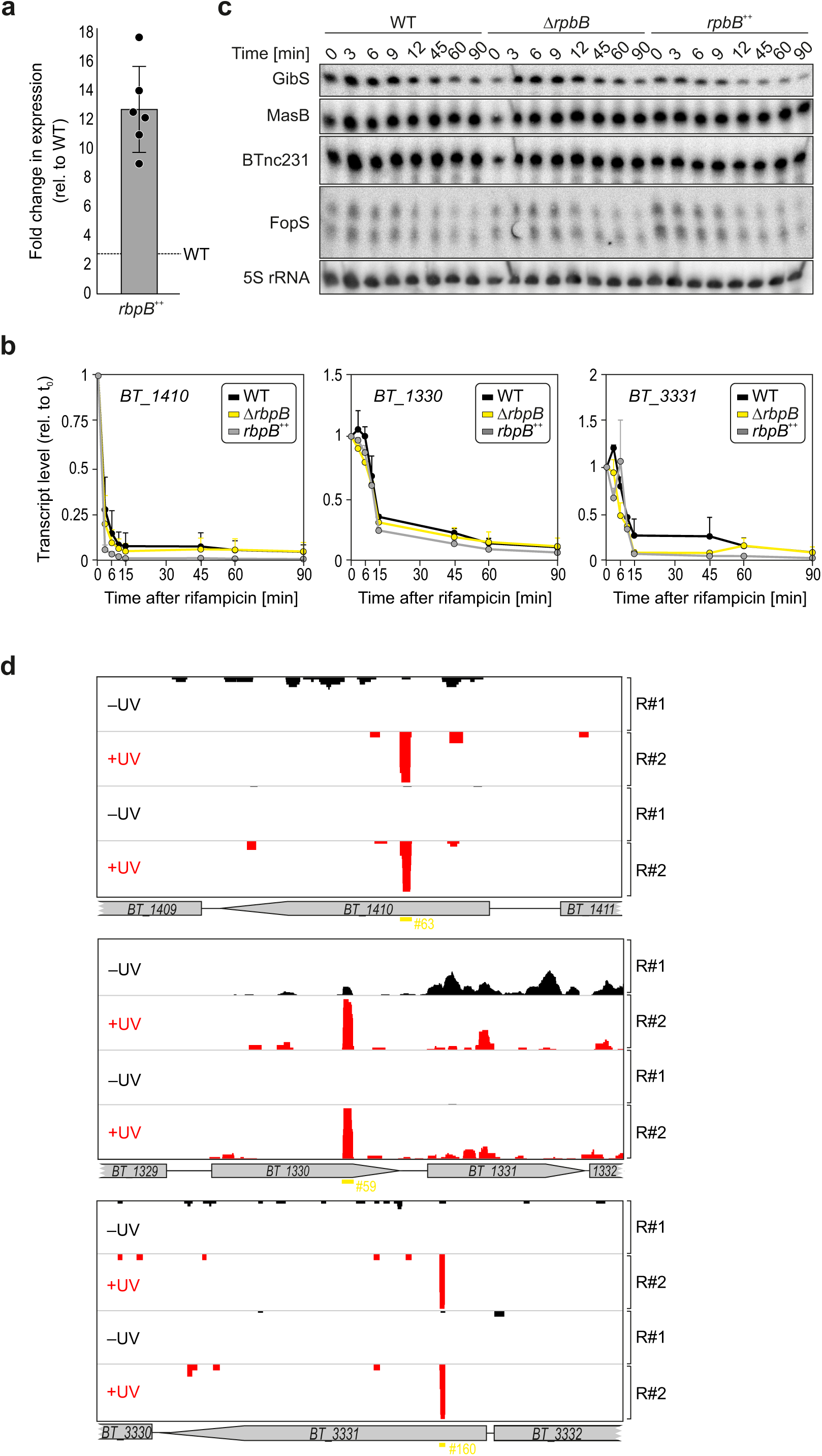
The cellular half-lives of selected mRNA ligands of RbpB depend on this protein’s concentration. **a,** Characterization of the *rbpB* overexpression strain (*rbpB* ^++^) used in this study. Plotted is the fold-change in expression of the *rbpB* mRNA in *rbpB* ^++^ *B*. *thetaiotaomicron* relative to its level in the isogenic wild-type as determined by qRT-PCR measurement and normalization against the 16S rRNA transcript (six biological replicates). Note, however, that strain *rbpB* ^++^ expresses a C-terminally FLAG-tagged variant of RbpB (as was used for CLIP). **b, c,** Rifampicin assay to measure the stability of RbpB-associated RNA ligands. Total RNA was collected from wild-type, Δ*rbpB*, and *rbpB* ^++^ *B*. *thetaiotaomicron* cultures grown in TYG to mid-exponential phase prior to rifampicin treatment (final concentration: 500 µg/mL) for the indicated time periods. The purified RNA samples were DNase-digested and subjected to qRT-PCR-based quantification of the decay kinetics of selected mRNA ligands of RbpB (top three mRNA interactors based on the enrichment score compared to the non-crosslinked control), normalized against 16S rRNA levels (**b**; plotted are the means +/−SD from three independent replicate measurements). Alternatively, RNA samples were loaded on a northern blot and probed with sequence-specific, radioactively labeled oligonucleotides against established sRNAs ^11,12^ that were significantly enriched in the RbpB CLIP-seq data, or a probe for 5S rRNA as a control (**c**; blots are representative of three biological replicates). **d,** Relative position of RbpB CLIP-seq peaks (peak ID marked in yellow) within the mRNA ligands, whose stability was quantified in panel b.

**Supplementary Figure S4:**
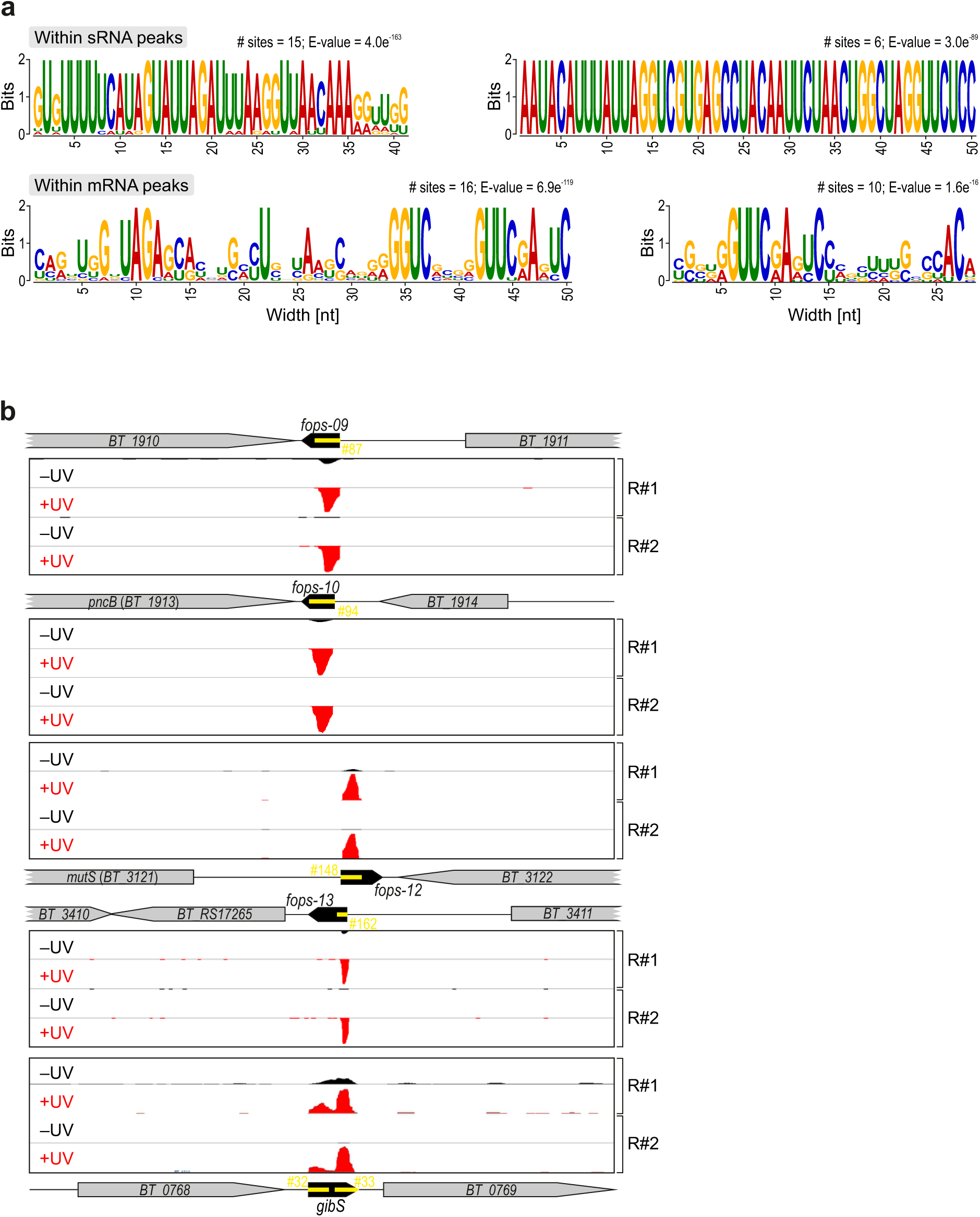
RpbB binds to specific sequence motifs. **a,** Most overrepresented sequence motifs in RbpB peak regions. MEME results for the top-enriched motifs in sRNAs (upper) and mRNAs (lower); their number of sites and E-value are given. **b,** Representative CLIP peaks within sRNAs harboring the 41-nt motif depicted in panel a (upper left). Shown are the scale-matched read coverages in control (black) and cross-linked (red) libraries. Annotations of coding sequences (grey arrows) and sRNA genes (black arrows) are given. Yellow horizontal lines and yellow font indicate the inferred peak positions and peak IDs, respectively.

**Supplementary Figure S5:**
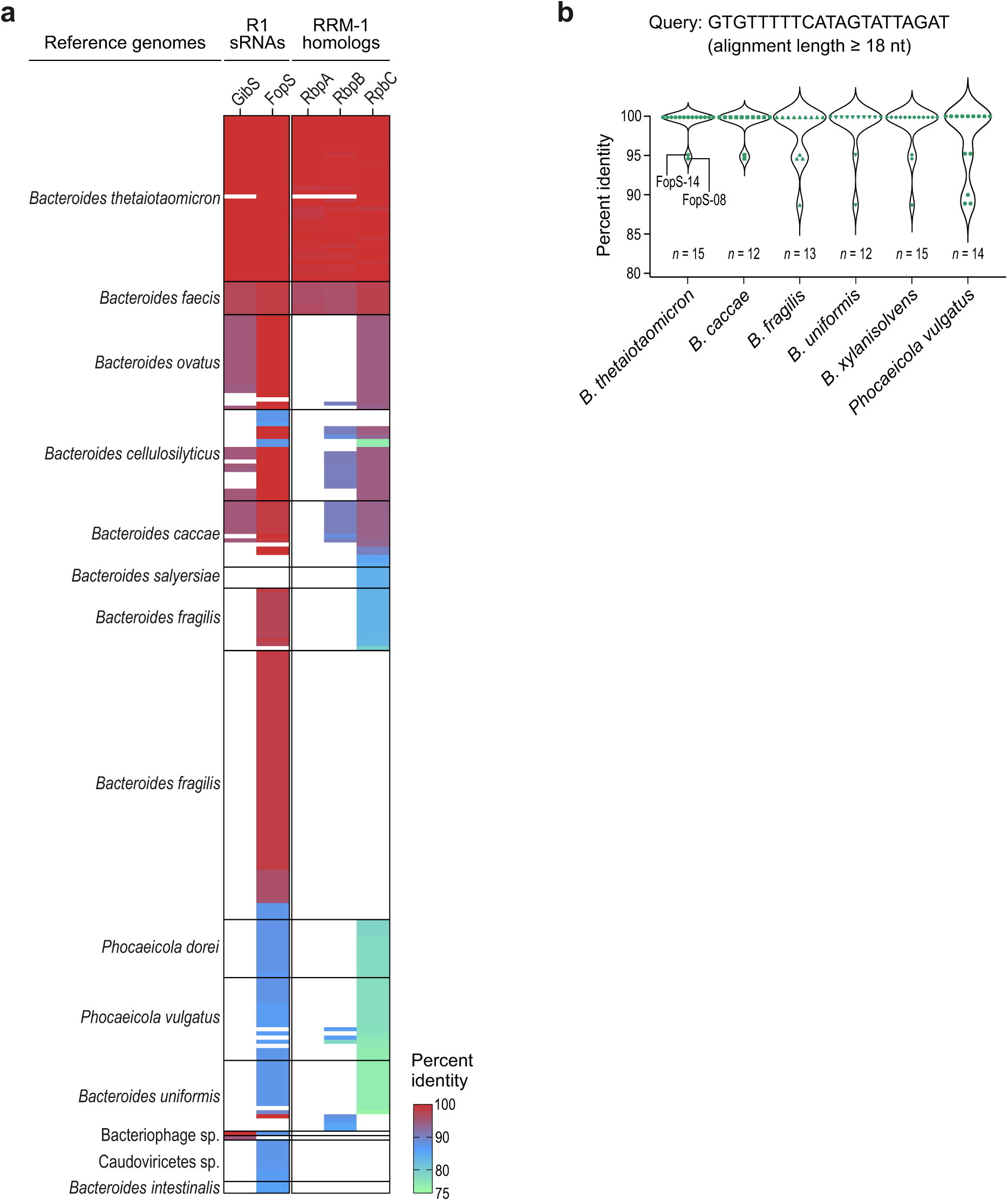
Conservation and prevalence of R1 sequence-containing sRNAs and of homologs of RRM-1 proteins across the *Bacteroidota*. **a,** BLAST analysis of GibS, the FopS family, and the three RRM-1 proteins RbpA, -B, and -C. In case of the FopS cluster, we blasted for FopS8, which—based on CD-HIT clustering ^85^ with 65% identity—is most representative for this sRNA family. The alignments were performed at default settings, except with increasing the maximum number of aligned sequences to 250. The heat map is color-coded based on sequence identity ranging from 75% (green) up to 100% (red). The Caudovirales phage was predicted to target *Bacteroides* based on matching CRISPR spacers^31^. **b,** Prevelance of R1 sequence copies within prominent *Bacteroidota* genomes, as inferred from a BLAST analysis.

**Supplementary Figure S6:**
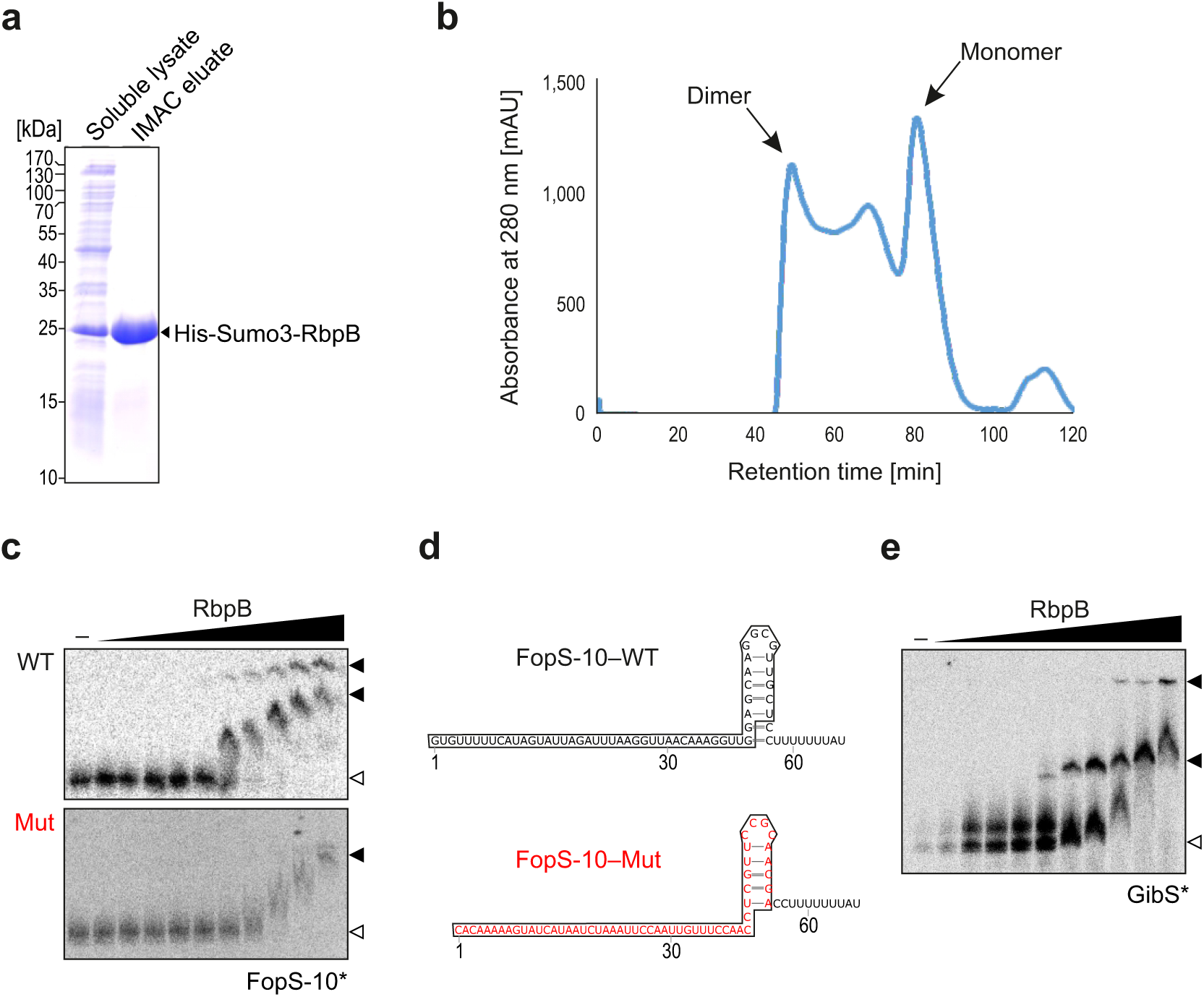
*In* -*vitro* validation of RbpB–sRNA interactions. **a,** Purification of recombinant His-Sumo3-RbpB expressed over night in *E*. *coli* Bl21 at 18°C. The soluble lysate was subjected to immobilized metal affinity chromatography (IMAC) and analyzed by SDS-PAGE. The tagged protein has a molecular weight of 23.4 kDa (RbpB itself has 11.0 kDa). **b,** Size-exclusion chromatography. Shown is the elution profile of RbpB, with two separate elution peaks at 68 and 80 mL, indicative of RbpB monomers and dimers. **c,** FopS-10 EMSAs. Increasing concentrations of purified RbpB (80 µM max.) were incubated with either wild-type T7-transcribed and 5’ end-labeled FopS-10 or a mutated variant thereof (4 nM). White and black arrows refer to free and bound FopS-10, respectively. **d,** Mutation of the RbpB binding site within FopS-10. Depicted is the sequence and predicted secondary structure of FopS-10 variants: wild-type (black) and mutated (red). The 55 nt-long RbpB-binding site is boxed. **e,** *In* - *vitro* -transcribed and 5’ end-labeled GibS (4 nM) was incubated with increasing concentrations of RbpB (80 µM maximum). White and black arrows refer to free and bound GibS, respectively.

**Supplementary Figure S7:**
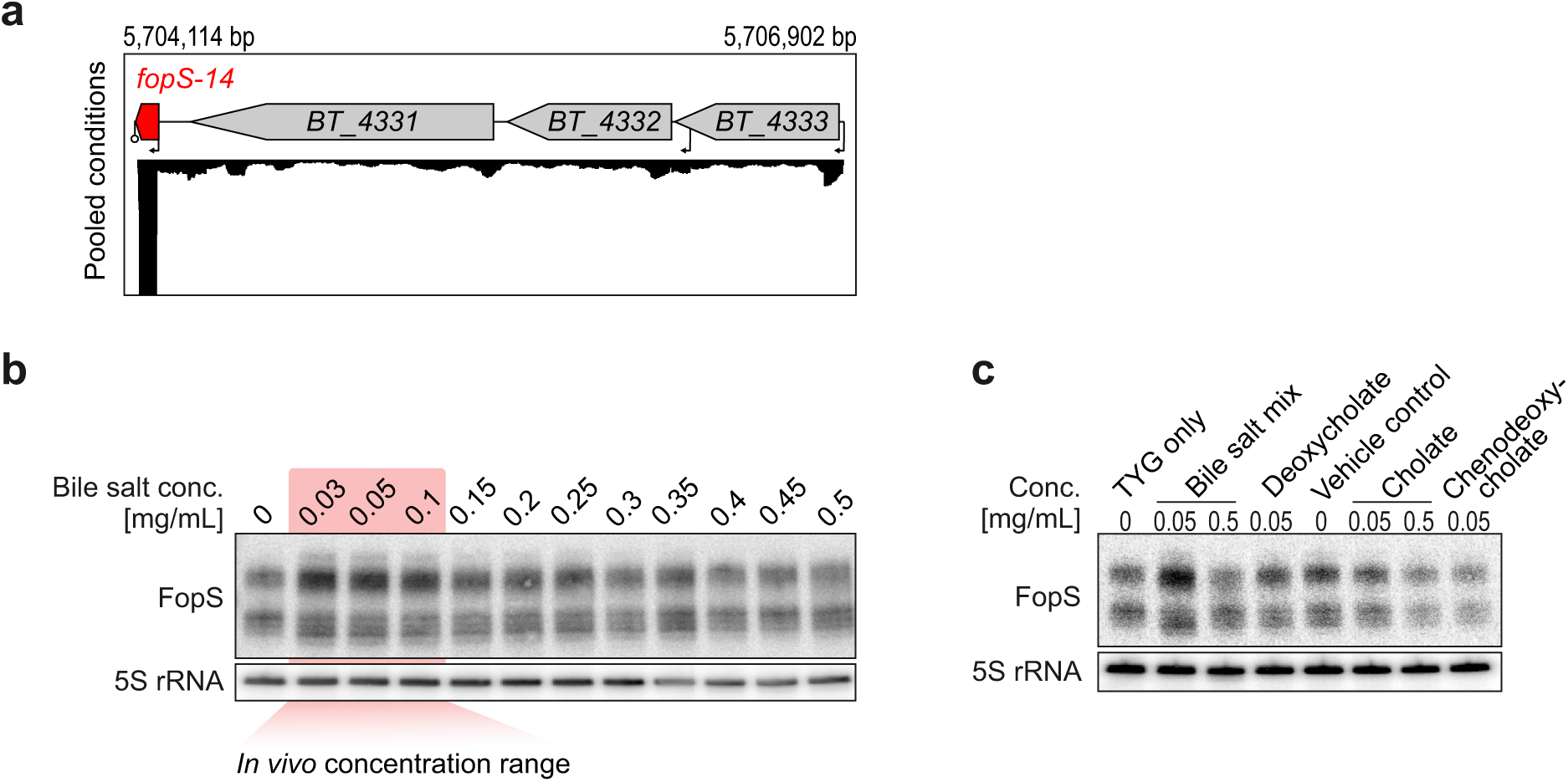
Expression profiling of FopS sRNAs. **a,** FopS-14 is a 3’-derived sRNA. The combined read coverages from Theta-Base ^12^ across the *fopS-14* locus is shown. **b, c,** Northern blot analysis of FopS sRNA expression across a bile salts concentration range, (0–0.5 mg/mL) (**b**) and upon exposure to the defined bile salts deoxycholate, cholate, and chenodeoxycholate, in the indicated concentrations (**c**).

**Supplementary Figure S8:**
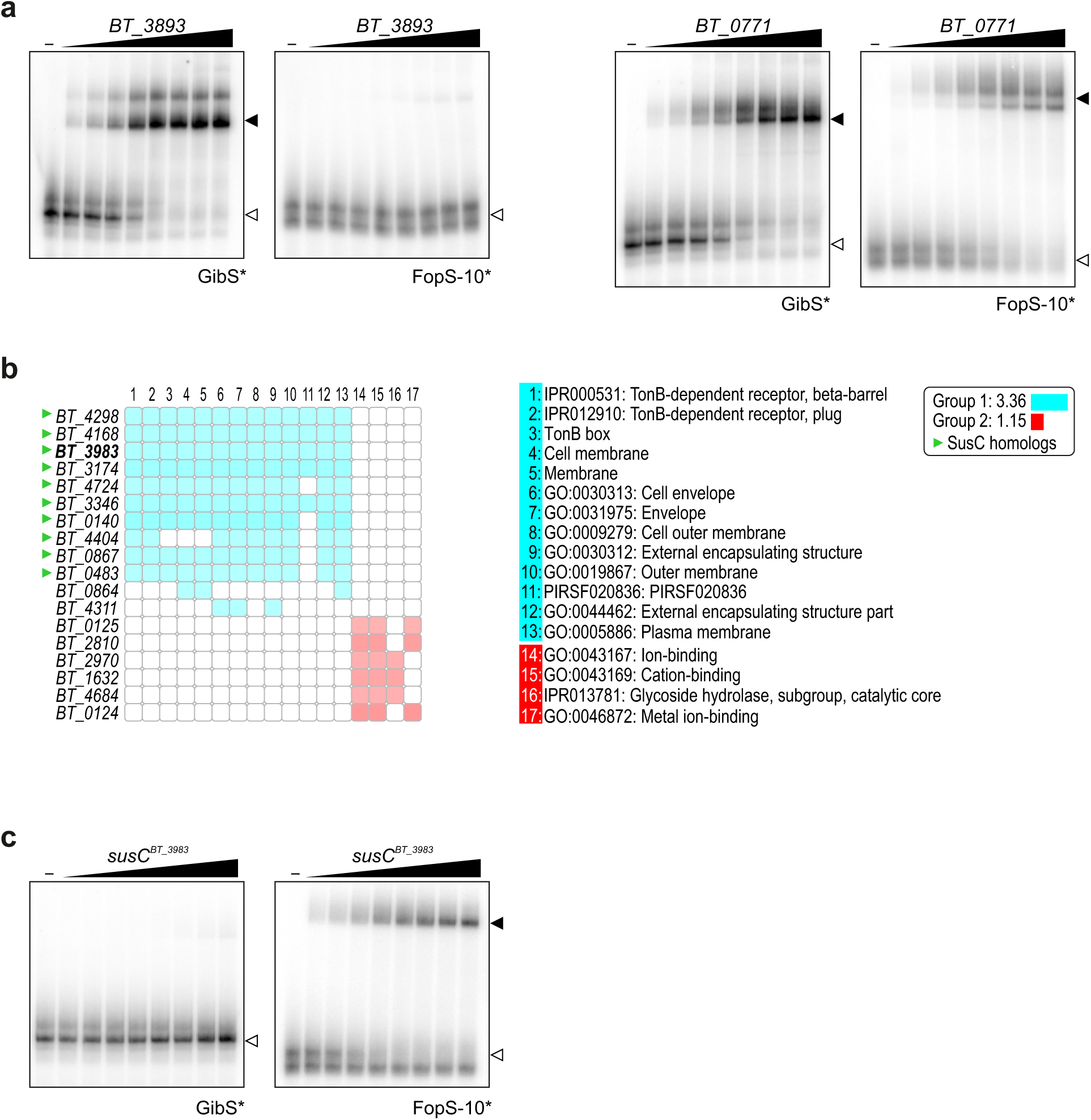
FopS target identification and validation. **a,** EMSAs with established GibS targets. *In* -*vitro* -transcribed and 5’ end-labeled GibS or FopS-10 (4 nM each) was incubated with increasing concentrations (up to 1,000 nM) of ∼150 nt long 5’ segments of either *BT_3893* or *BT_0771*. White or black arrows refer to free or bound GibS and FopS-10, respectively. **b,** Functional classification of the top 18 putative FopS targets as predicted by IntaRNA revealed two large clusters (blue and red squares) with a significant enrichment of SusC homologs (green arrowheads). **c,** EMSAs to confirm FopS binding to the predicted target mRNA *susC^BT_3983^*. Experiment analogously performed to the one described in panel a, except that a ∼180 nt long 5’ segment of susC^BT_3983^ was used as a target fragment.

**Supplementary Figure S9:**
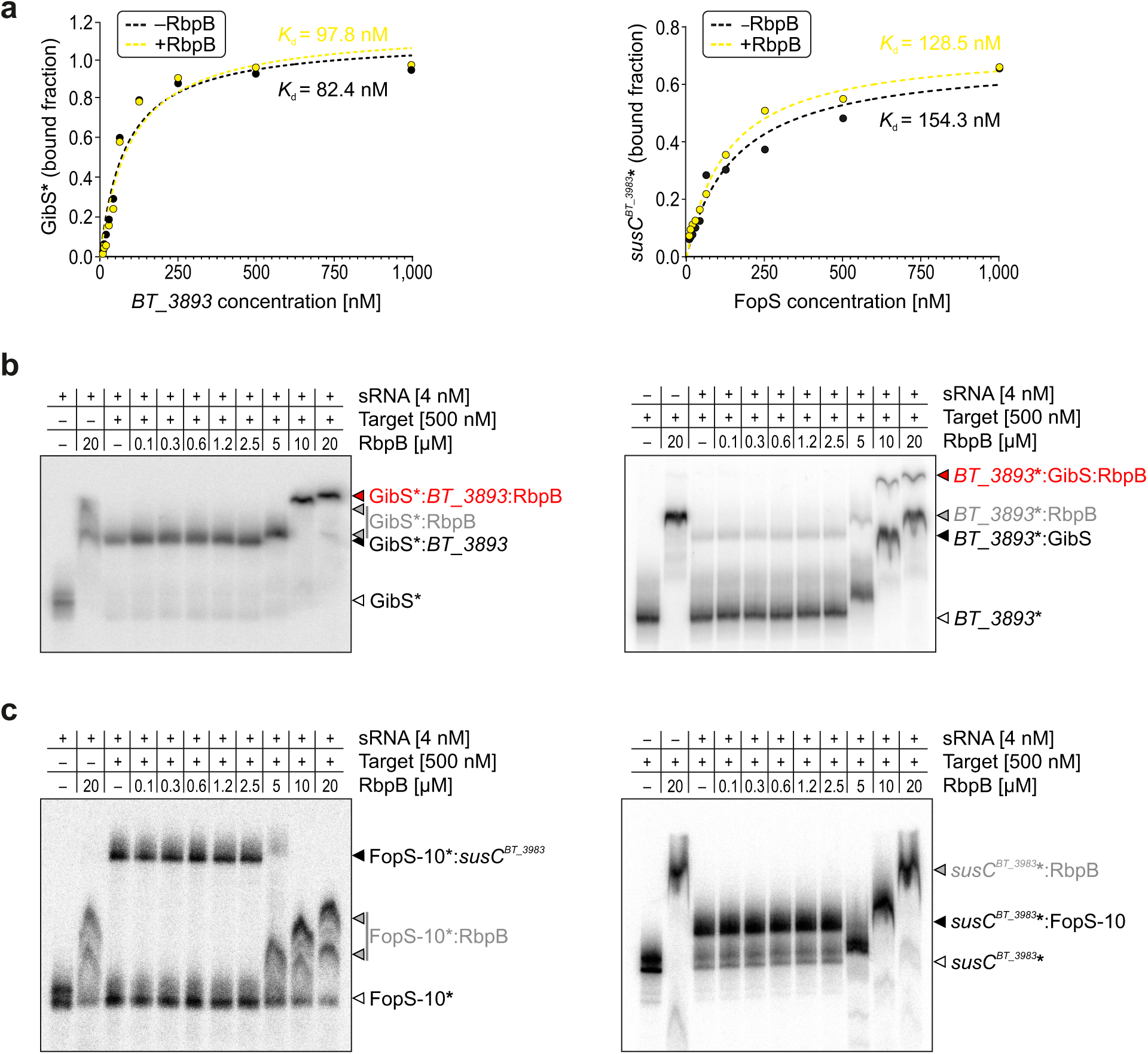
*In* -*vitro* interaction of GibS and FopS with their targets in the presence or absence of RbpB. **a,** Quantification of EMSAs. K*_d_* values represent the means of three independent replicate experiments performed with either *in*-*vitro* -transcribed and 5’ end-labeled GibS or a ∼180 nt long 5’ region of *susC^BT_3983^* (4 nM each) incubated with increasing concentrations of a 5’ segment of *BT_3893* (∼150 nt) or FopS-10, respectively. EMSAs were performed in the absence (black line) or presence (yellow line) of 1 µM RbpB. **b,** EMSA reveals the 5’ region of *BT_3893* mRNA to be bound by RbpB *in vitro*. In the presence of the GibS sRNA, a trimeric complex formed. **c,** Three-component EMSA analogous to panel b, but with FopS-10 and its target mRNA, *susC^BT_3983^*.

**Supplementary Figure S10:**
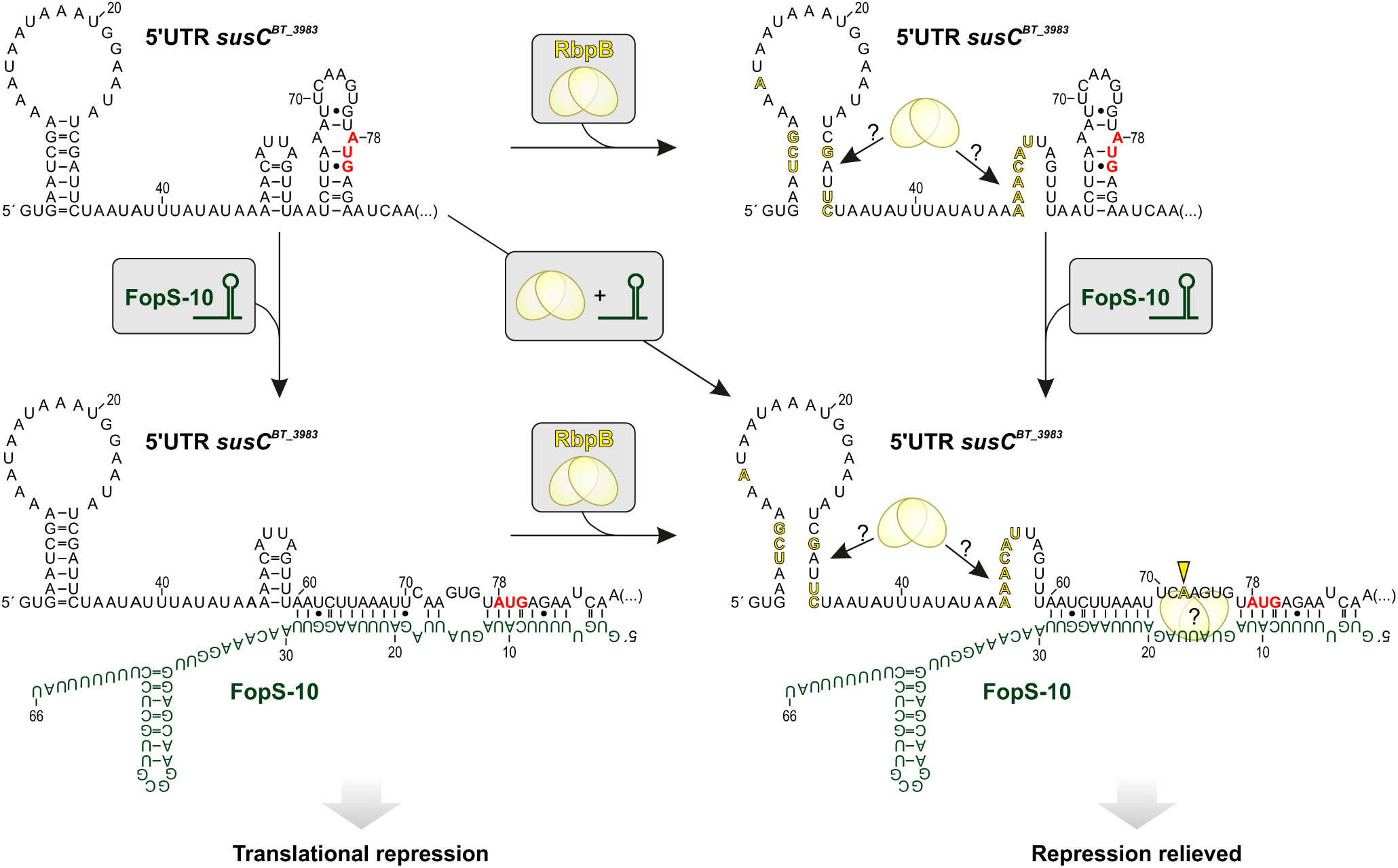
Modelling the interaction of the 5’ portion of *susC ^BT_3893^* with FopS-10, in the presence or absence of RbpB. Intra- and inter-molecular base-pair interactions were inferred from the band pattern in the inline probing assay (Fig. 4e). Ribobases involved in rearrangements upon RbpB association (i.e., residues with an altered cleavage efficiency; yellow arrowheads in Fig. 4e) are depicted in yellow font and the translational start codon of the *susC^BT_3893^* mRNA in red font. The yellow arrowhead in the lower right schematic highlights the structural change at position −6 relative to the translational start site that occurs specifically only when both, FopS-10 and RbpB, are present (enhanced cleavage at position 72 in Fig. 4e). The exact binding site of RbpB in the 5’ single-stranded region of FopS-10 was inferred from CLIP-seq (see Suppl. Fig. S4b). Since *susC^BT_3893^* was not expressed in the CLIP-seq condition, the precise binding site of RbpB in this mRNA is unknown; however, the evoked changes in the cleavage pattern in the presence of RbpB could imply that protein binding melts the two 5’-most stem loops (as indicated in the model by arrows with question marks).s

**Supplementary Table 1: Combined CLIP-seq data.**

**Supplementary Table 2: List of bacterial strains, plasmids, and oligonucleotides used in this study.**

## ACKNOWLEDGEMENT

We would like to acknowledge the recombinant protein expression facility of the Rudolf-Virchow-Center (University of Würzburg) for expression and purification of the RbpB protein. We thank Milan Gerovac and members of the Westermann and Faber labs for fruitful discussions of this work. We are grateful to Anke Sparmann, Jörg Vogel, Chase Beisel, Lena Amend, and Thomas Guest for critical reading and constructive feedback on this manuscript. This work was funded by the German Research Foundation (DFG; Individual Research Grant We6689/1-1). Research in the Westermann laboratory is supported by the European Research Council (ERC Starting Grant #101040214).

